# Influence of root cortical aerenchyma on the rhizosphere microbiome of field-grown maize

**DOI:** 10.1101/2023.01.31.525837

**Authors:** Tania Galindo-Castañeda, Claudia Rojas, Ulas Karaöz, Eoin L. Brodie, Kathleen M. Brown, Jonathan P. Lynch

## Abstract

The root anatomical phenotype root cortical aerenchyma (RCA) decreases the metabolic cost of soil exploration and improves plant growth under drought and low soil fertility. RCA may also change the microenvironment of rhizosphere microorganisms by increasing oxygen availability or by reducing carbon rhizodeposition. We tested the hypothesis that plants with contrasting expression of RCA have different rhizosphere prokaryotic communities. Maize inbreds were grown in two field sites, Limpopo Province, South Africa and Pennsylvania, USA, and their rhizosphere soil sampled at flowering. High- and low-nitrogen fertilization was imposed as separate treatments in the experiment in South Africa. The rhizosphere microbial composition of plants with contrasting RCA was characterized by metabarcoding of the 16S rRNA genes. Geographic location was the most important factor related to the composition of rhizosphere microbial communities. In the site in South Africa, RCA explained greater percent of variance (9%) in the composition of microbial communities than genotype (7%). Although other root anatomical and architectural phenotypes were studied as possible cofactors affecting the microbial composition, RCA was among the best significant explanatory variables for the South African site although it was neutral in the Pennsylvania site. High-RCA rhizospheres significantly enriched OTUs of the families *Burkholderiaceae* (in South Africa) and *Bacillaceae* (in USA), compared to low-RCA plants, and OTUs of the families *Beijerinckiaceae* and *Sphingomonadaceae* were enriched at the two nitrogen levels in high RCA plants in South Africa. Our results are consistent with the hypothesis that RCA is an important factor for rhizosphere microbial communities, especially under suboptimal nitrogen conditions.

## INTRODUCTION

Root-associated microbes alter plant nutrition and plant health, becoming a key aspect of root biology for the development of sustainable agriculture. Factors such as soil type, geographic location, agronomic practices, plant taxa, plant age, and root exudates, are modifiers of rhizosphere microbial communities as demonstrated by metagenomic analyses (reviews by Compant et al. 2019 and Philippot et al. 2013). Despite the fact that roots create and structure the niches of rhizosphere communities, the effects of specific root phenotypes on microbial communities are less well known. Promising root phenotypes that improve soil resource acquisition can be targeted by plant breeding programs to produce new cultivars suited for the challenges of modern agriculture (Lynch 2019). However, selection for such phenotypes should not compromise beneficial microbial associations in the rhizosphere in order to meet the requirements of sustainable crop production. This is especially true for nitrogen, since microbial transformations play key roles in regulating nitrogen availability in the rhizosphere by means of nitrogen fixation (Dobbelaere et al. 2003), ammonification, nitrification and denitrification (Hai et al. 2009; Hinsinger et al. 2009; Li et al. 2014; Zhao et al. 2017). Therefore, understanding the effects of promising root phenotypes on microbial communities under nitrogen limitation is an important element of breeding crops with reduced requirements for nitrogen fertilizer.

Nitrogen fertilization is a primary economic, environmental, and energy cost of intensive maize production (FAO 2017; Robertson and Vitousek 2009). Only 33% of the nitrogen applied to cereal crops is recovered as grain, the rest, which remains as vegetative biomass or is lost to the environment, accounts for approximately $15.9 billion annual loss (Raun and Johnson 1999). A complex microbial network participates in the nitrogen cycle in the rhizosphere (Højberg et al. 1996; Neumann and Römheld 2012; Van Deynze et al. 2018), with the predominant processes depending on the microhabitats created by roots, which we propose, are in large measure determined by the root architecture and anatomy.

One strategy to help ameliorate excessive use of nitrogen fertilizers and the loss of nitrogen in maize fields is the selection of cultivars with specific root phene states (phenes are the elements composing a phenotype (York et al. 2013)) that improve nitrogen capture (Gaudin et al. 2011; Lammerts van Bueren and Struik 2017; Lynch 2013; Lynch 2015; Lynch 2019). Specifically, increased root cortical aerenchyma (RCA), the production of air pockets in the cortical tissue, has been reported as a response of maize to low nitrogen stress (York et al. 2015). In addition, plants with increased RCA had increased grain yield under suboptimal nitrogen fertilization (Saengwilai et al. 2014). These results are in accord with the concept that increased RCA reduces the metabolic cost of soil exploration (Lynch, 2015) especially under resource scarcity, by means of reduction of metabolically active tissue in the root cortex, and indicate that plant breeding programs could deploy RCA to develop maize cultivars better adapted to low-nitrogen stress (Lynch 2019).

Root anatomical phenotypes may change the conditions of niches inhabited by rhizosphere microbes. The production of RCA has major impacts on the rhizosphere by changing oxygen availability and shifting facultative or anaerobic microbial functions towards aerobic metabolisms in the vicinities of RCA air pockets (Arth and Frenzel 2000; Li et al. 2008; Risgaard-Petersen and Jensen 1997). Processes like carbon utilization, nitrogen transformation, and metal accumulation depend on soil oxygen content and redox potential (Neumann and Römheld, 2012), and may modify microbial communities in the rhizosphere. Likewise, roots with reduced RCA may restrict oxygen diffusion to the rhizosphere, thereby limiting aerobic microbial metabolism and nutrient utilization.

Previous studies have analyzed the microbial composition of the rhizosphere using high-throughput amplicon sequencing of soils associated with agriculturally relevant plants, including maize (Bakker et al. 2015; Dohrmann et al. 2013; Li et al. 2014; Peiffer et al. 2013, Walters et al. 2018). Root architectural traits sus as root class and order (reviewed by Saleem, 2018), and specific root length are significant factors explaining microbial community composition of the rhizosphere (Pérez-Jaramillo et al. 2017). Also, decrease in specialist microbial OTUs were reported from finer to coarser root classes of two cultivars of field-grown *Nicotiana tabacum* (Saleem et al. 2016). To our knowledge, no high-throughput amplicon sequencing studies of the rhizosphere have studied RCA and its associations with rhizosphere microbial communities under nitrogen stress. The study of phenotypic effects on the rhizosphere microbiome under low nutrient stress is important to inform plant-breeding programs targeting root phenes and better root microbiomes in the context of sustainable agriculture. This study addressed the effects of RCA on the composition of rhizosphere bacterial and archeal communities (prokaryotes, referred as microbial communities here on) of maize under field conditions, focusing on the comparison of specific OTUs between plants with contrasting levels of RCA under optimal and suboptimal nitrogen fertilization. Using metabarcoding of 16S rRNA genes of total rhizosphere DNA, combined with root phenotyping in two experimental maize fields, we tested the hypothesis that RCA has significant effects on the microbial community composition.

## MATERIALS AND METHODS

### Field experiment and sampling

#### Experimental conditions and plant material

Two experiments were conducted, one at the Russell E. Larson Research and Education Center of the Pennsylvania State University in Rocksprings, PA, USA (designated herein as **RS**) (40°42′37″.52 N, 77°57′07″.54 W, 366 masl), from June – August 2012; the other at The Ukulima Root Biology Research Center (designated herein as **URBC**), Limpopo province, Republic of South Africa (24°33′00.12 S, 28°07′25.84 E, 1235 masl) from December 2013 to February 2014. Recombinant inbred maize lines (RILs) differing in RCA formation from the IBM population (B73 x Mo17) (URBC High-RCA: IBM031, IBM196; URBC Low-RCA: IBM001, IBM345; RS High-RCA: IBM031, IBM034, IBM177, RS-Low-RCA: IBM001, IBM157, IBM338) (Kaeppler et al. 2000; Senior et al. 1996) were planted in three row-plots with 0.76 m inter-row spacing and 0.23 m in-row spacing for a final population of 57,278 plants*ha^-1^. The soil at the experimental sites consisted of a Hagerstown silt loam (fine, mixed, semiactive, mesic Typic Hapludalf) at RS and a clovelly loamy sand (Typic Ustipsamment) at URBC. Soil test reports from the two sites are summarized in Supplementary Table 1. Contrasting levels of nitrogen fertilization were imposed at URBC according to soil analyses at the beginning of the field season in order to provide low nitrogen (33 kg*ha^-1^ applied at URBC) treatment to half of the blocks and high nitrogen conditions (fertilized with 207 kg*ha^-1^ at URBC, and 150 kg*ha^-1^ at RS) to the other half. At RS, each block was a 0.4 ha separate field and at URBC the blocks were randomly distributed in a 20-ha irrigation pivot. In both locations, all nutrients except nitrogen were adjusted to meet the requirements for maize production as determined by soil tests. Pest control and irrigation were carried out as needed. The RS experiment was a complete randomized block design and the URBC experiment was a completely randomized design with genotypes as treatments. The experiment at RS had three replicates, and the experiment at URBC had four replicates; all replicates were designated as blocks.

#### Rhizosphere and bulk soil sampling

Two plants per genotype were excavated from the central row at flowering (12 weeks after planting at RS and 13 weeks after planting at URBC) with a shovel inserted approximately 40 cm radial distance from the stem, and 30 cm depth. The root systems were processed similar to Lundberg et al. (2012) with modifications in order to scale the method to maize and field studies. Briefly, the root crowns were excavated, kept in paper bags and immediately transported to the sample processing station, adjacent to the field. The root crowns were carefully shaken and ten nodal roots (two to three root segments per whorl, from the second to the fifth whorl) per plant were aseptically clipped and placed into 250 mL sterile plastic bags. A total of 20 root fragments (∼40 g fresh weight) per plot were collected. The samples were kept at 4°C for maximum 24 h, and the rhizosphere samples collected in 150 mL of a 20% sterile Tween®20 solution (Amresco, Inc., Solon, OH, USA) poured into the plastic bag. Each closed plastic bag containing roots, soil and tween solution was manually shaken for 1 min. Then, the solution with the released soil was filtered with nylon cell strainers (MACS® SmartStrainers, 100 µm), and the filtrate centrifuged at 3,000 g for 15 min. The soil pellet was immediately processed for DNA extraction for RS, and stored at −20°C for 24 h and then placed at −70°C for URBC. For the URBC samples, the frozen soil pellet was lyophilized (Labconco System Freezone, 1L freeze-drier coupled to a 117 L*min^-1^ vacuum pump) for 48 h to constant weight. Three samples of bulk soil were taken per plot using a corer of 5.1 cm diameter inserted 20 cm depth in locations free of plant roots in the furrow, pooled (all the samples coming from the different genotypes were collected in the same field replicate), and ∼ 10 g diluted in tween solution and filtered through nylon cell strainers and the filtrate processed as described for rhizosphere samples. The lyophilized soil samples were aseptically stored in 2 ml vials at 4°C for 2 weeks and transported to the USA for DNA processing.

#### Sampling for root phenotyping

Three root crowns per plot were excavated and sampled for anatomical analysis as previously described (York et al. 2015). These plants were different to the plants used for DNA extraction with the purpose of avoiding changes in root anatomy due to the DNA extraction processing on the roots used for rhizosphere soil collection. Root anatomy was measured on root cross-sections with the software *RootScan* (Burton et al. 2012). At URBC, two of the three plants selected for anatomical sampling were also used for architecture phenotyping with “DIRT” (Bucksch et al., 2014). Washed root crowns were imaged on a table with black background using a Nikon D70s digital camera with focal length ranging 22 – 29 mm, exposure time of 1/30 – 1/50 sec., maximal aperture of 3.6 – 4.1, and digital zoom only. The camera was mounted on a tripod at 50 cm above the imaging board. All the images were taken at a resolution of 3,008 x 2,000 pixels. For RCA, individual values were assigned to qualitative ranks (high, intermediate and low) based on values of percentage of cortical area that is aerenchyma in order to facilitate some statistical analysis (PERMANOVAS to compare genotype vs. phenotype effect, and to plot PCoAs by RCA levels) as described below (see Fig. 4).

### DNA extraction

Soil samples weighing 0.25 g of either of centrifuged rhizosphere (at RS) or lyophilized soil (at URBC) were processed with the PowerLyzer Power Soil DNA Kit extraction (MoBio Laboratories, Inc., Carlsbad, CA). Concentration and quality (260/280 and 230/260 absorbance ratios) were measured with a NanoDrop 1000 Spectrophotometer (Thermo Fisher Scientific Inc.). Integrity of the extracted DNA was confirmed (> 10 kb) in 0.8% agarose electrophoresis gels (110V for 1.5 h) by comparison of the extracted DNA stained with ethidium bromide with a molecular weight marker (2-Log DNA ladder, New England Biolabs® Inc.). The DNA samples were stored at −70 °C (18 months for RS and a week for URBC). Double-stranded DNA concentration was measured by fluorescence with a SPECTRAmax GEMINI-XPS microplate reader (Molecular devices, Sunnyvale, CA, USA) and with picogreen nucleic acid stain.

### 16S rRNA amplification and sequencing

DNA concentrations were normalized to 1 – 5 ng µl^-1^, and used for triplicate PCRs with the 515F-806R primer pair, targeting archaeal and bacterial 16S rRNA, including barcodes as previously described (Caporaso et al. 2012). PCR conditions and product purification were as follows: one denaturation cycle at 94°C for 3 min followed by 30 annealing cycles (95°C for 30 sec, 52°C for 45 sec, and 72°C for 90 sec); and an extension cycle at 72°C for 12 min, and hold at 4°C in a MyCycler thermal cycler (Bio-Rad, Hercules, CA, USA). PCR products of the three reactions were pooled into a single sample, purified with 0.1% carboxyl-modified Sera-Mag Magnetic Speed-beads^TM^ (Fisher), and eluted in 1x TE for quantification. Concentrations of the purified PCR products of individual samples were determined by fluorescence with a SPECTRAmax GEMINI-XPS microplate reader (Molecular devices, Sunnyvale, CA, USA). The quality was assessed via a 2100 Bionalyzer (Agilent Technologies, Wilmington, USA). The samples were then pooled into a 25 ng*µL^-1^ (57.7 nM) library. Number of Illumina-amplifiable DNA fragments in the library were 2nM, determined by qPCR with the KAPA kit (Biosystems. Boston, MA, USA) and confirmed with a 2-point Qubit 2.0 fluorometer (Invitrogen). The library was denatured with 10 µL of 0.2N NaOH and diluted in a solution of denatured PhiX, for a final library concentration of 10 pM. Sequencing of the amplicons were performed in an Illumina MiSeq system (Illumina, San Diego, CA, USA) with 500 cycles.

### Sequence analysis

The Illumina sequence data was demultiplexed, quality filters applied, dereplicated, and OTUs (Operational Taxonomic Units) assigned as previously described with the pipeline UPARSE with default options: Quality score of 16, OTU radius of 3%, and no length trimming (Edgar 2013). We used 97% similarity cutoff to assign biological identities to the OTUs by comparison against the database SILVA (Quast et al. 2013). Non-classified OTUs at the domain levels (Bacteria or Archaea), chimera and singleton sequences were discarded with Qiime (Caporaso et al. 2010). Dataset preparation for downstream analysis including the taxonomy, OTU count table, phylogenetic tree and sample information was performed with the R package Phyloseq (McMurdie and Holmes 2013; R Core Team, 2020).

### Data analysis

#### OTU preprocessing

The obtained OTUs were analyzed separately by experiment. Low-count OTUs detected less than more than 2 times in fewer than at least 10% of the samples were eliminated from the OTU table of each experiment.

#### Species diversity

For diversity calculations one of the URBC samples with relatively low read-count (less than 10% of the second lowest read-count) and one sample with low OTUs (236 compared to 1115 OTUs in the second lowest OTU counts) at RS were dropped for further analyses. Alpha diversity analyses with the Shannon diversity index were performed on observed OTU counts. OTU tables were randomly subsampled without replacement (rarefied to the minimum number of OTUs for the samples of each experiments, 21268 for URBC and 22358 for RS) in order to perform beta diversity analyses. To study the beta diversity among samples, meaning the difference of shared microbial OTUs per taxa among different samples, we used UniFrac distance metrics, which measures the relatedness of samples based on phylogenetic distance of their taxa (Lozupone et al. 2010). Unweighted UniFrac distance takes into account the composition of each sample, while the weighted UniFrac accounts for the abundance of each taxa. Weighted and unweighted UniFrac distances on the rarefied OTU tables were used to perform principal coordinate analysis (PCoA), permutational multivariate analyses of variance (PERMANOVA), and constrained correspondence analyses (CCA). PCoA was used to observe patterns in sample aggregation by nitrogen levels at URBC, and by genotypes, rhizosphere versus bulk soil, and significant root phenotypes at the two sites. PERMANOVAS revealed the effect of nitrogen, genotype and RCA (transformed to qualitative ranges) on the microbial communities, and CCA was used to measure the variation in rhizosphere microbial communities explained by specific root phenes measured as a mean to understand the relative contribution of RCA in comparison with other root phenes. For CCA analysis we used quantitative values resulting from phenotyping. Due to the high number of phenotypic variables resulting from *RootScan* and DIRT, prior to the CCA, we performed a selection of the most significant variables with random permutations using the function ordistep of the R package Vegan (Oksanen et al. 2017). The resulting variables were then included in the model of the ordination. The significance of the model and of the phenes included in the model were calculated with permutation tests with the function anova.cca of the R package Vegan.

#### Phenotype-sensitive OTUs

Association of significantly enriched OTUs with contrasting root phenotypes were performed by fitting a generalized linear model with a negative binomial distribution to normalized abundance values. We used the “trimmed means of M” method for the normalization available through the BioConductor package edgeR (McCarthy et al. 2012; Robinson et al. 2010) and expressed the normalized counts as relative abundance counts per millions (CPM) for each OTU in each site and nitrogen level. To test for differential abundance, we used a likelihood ratio tests (LRT) with the R package edgeR. OTUs that were significantly increased or depleted (compared to the control treatment by a significantly different fold-factor, see results section for more details on the specific control treatment used for each comparison) with *P* values < 0.01, were considered phenotype-responsive.

#### Plant phenotyping

The effect of root architectural and anatomical phene states (with special focus on RCA) on UniFrac distance metrics was assessed with the scale-transformed quantitative values of the measured phenotypes retrieved by *RootScan* and DIRT. RCA was grouped into categorical states and assigned to each root sample for the PCoA.

## RESULTS

### Taxonomy and diversity

A total of 3,403,932 high quality sequences were obtained from the two experimental sites with a median read count per sample of 59,718 (range of 1,865 – 126,242). 17,693 microbial OTUs resulted from the alignment of the sequences with the SILVA dataset (Quast et al. 2013). After low-count OTU removal, the total sequences decreased from 951,299 to 941,051 at RS, and from 2,452,633 to 2,394,705 at URBC; and the total number of OTUs from 12,478 to 6,125 at RS and from 14,073 to 5,536 at URBC.

There were 34 and 45 phyla found at URBC and RS respectively. *Proteobacteria*, *Acidobacteria*, *Actinobacteria*, *Verrucomicrobia*, *Bacteroidetes*, and *Firmicutes* were among the most abundant phyla overall (Fig. 1). Rhizosphere soil at the two sites had a greater proportion of *Proteobacteria* and *Bacteroidetes*, and smaller proportion of *Acidobacteria* and *Nitrospirae* than bulk soil. *Gemmatimonadetes*, *Chloroflexi* and *Thaumarchaeota* were reduced in the rhizosphere at URBC but not at RS. Eleven phyla were unique to RS (for example, *Hydrogenedentes*, *Zixibacteria*, *Nitrospinae, Atribacteria*) while all the phyla found in URBC were also present in RS. *Sphingomonadaceae* and *Burkholderiaceae* (both from Phylum *Proteobacteria*) and *Micrococcaceae* (phylum *Actinobacteria*) were the most abundant families at URBC (Supplementary Fig. 1a). *Sphingobacteriaceae*, *Beijerinckiaceae*, and *Rhodanobacteraceae* were among the most abundant families at URBC but not among the most abundant families at RS. *Sphingomonadaceae, Burkholderiaceae,* and *Xanthobacteraceae* (Phylum *Proteobacteria*) were the most abundant families at RS (Supplementary Fig. 1b). RS had greater OTU richness compared to URBC, and bulk soil had greater average species diversity than rhizosphere soil at URBC (Fig. 2).

**Fig. 1.**
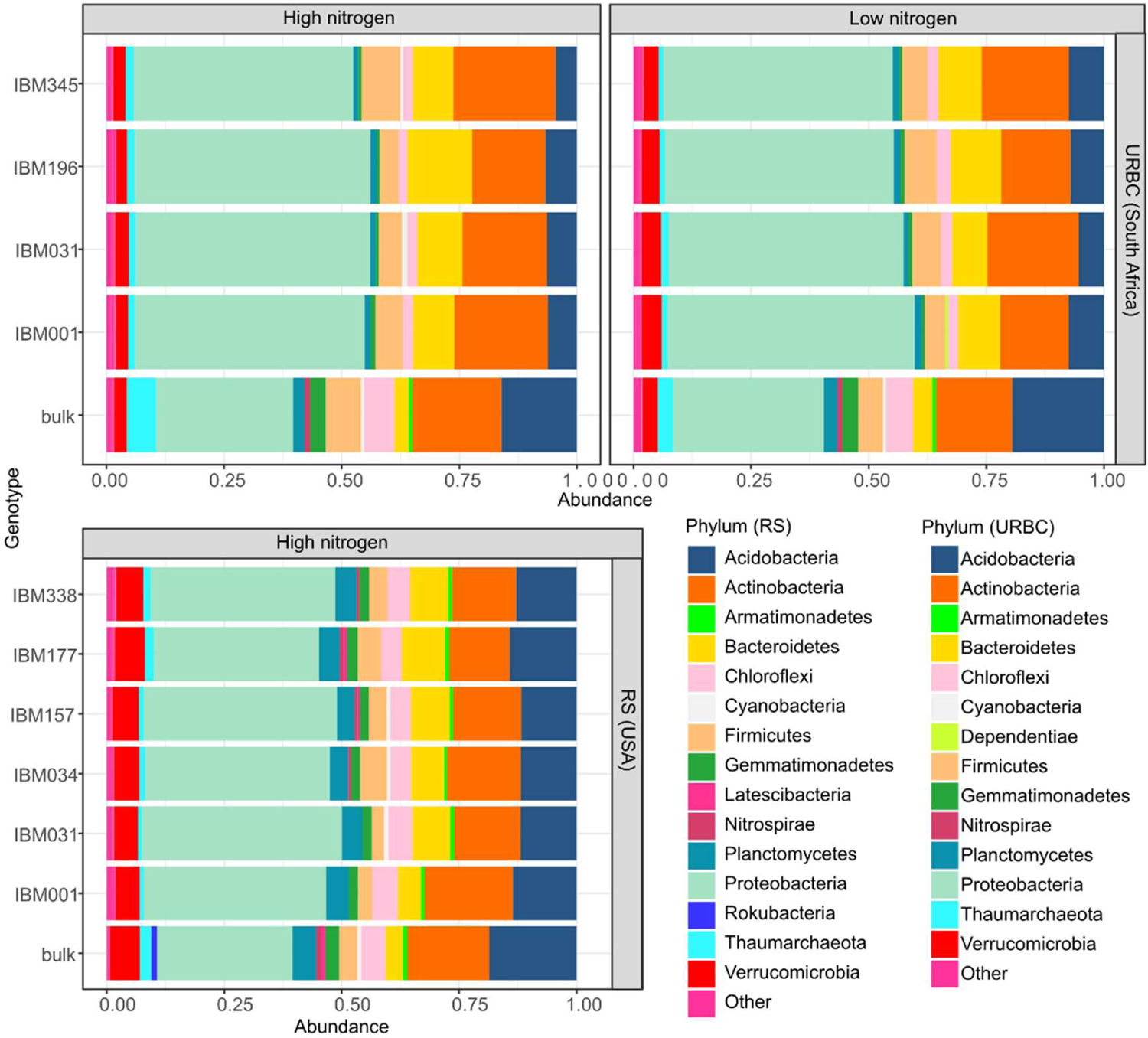
Bar plots of the relative abundances discriminating the 15 (for RS) and 14 (for URBC) most abundant phyla in each rhizosphere sample and bulk soil by experimental site. Low abundance phyla are represented as “other”. Values are means of at least three replicates.

**Fig. 2.**
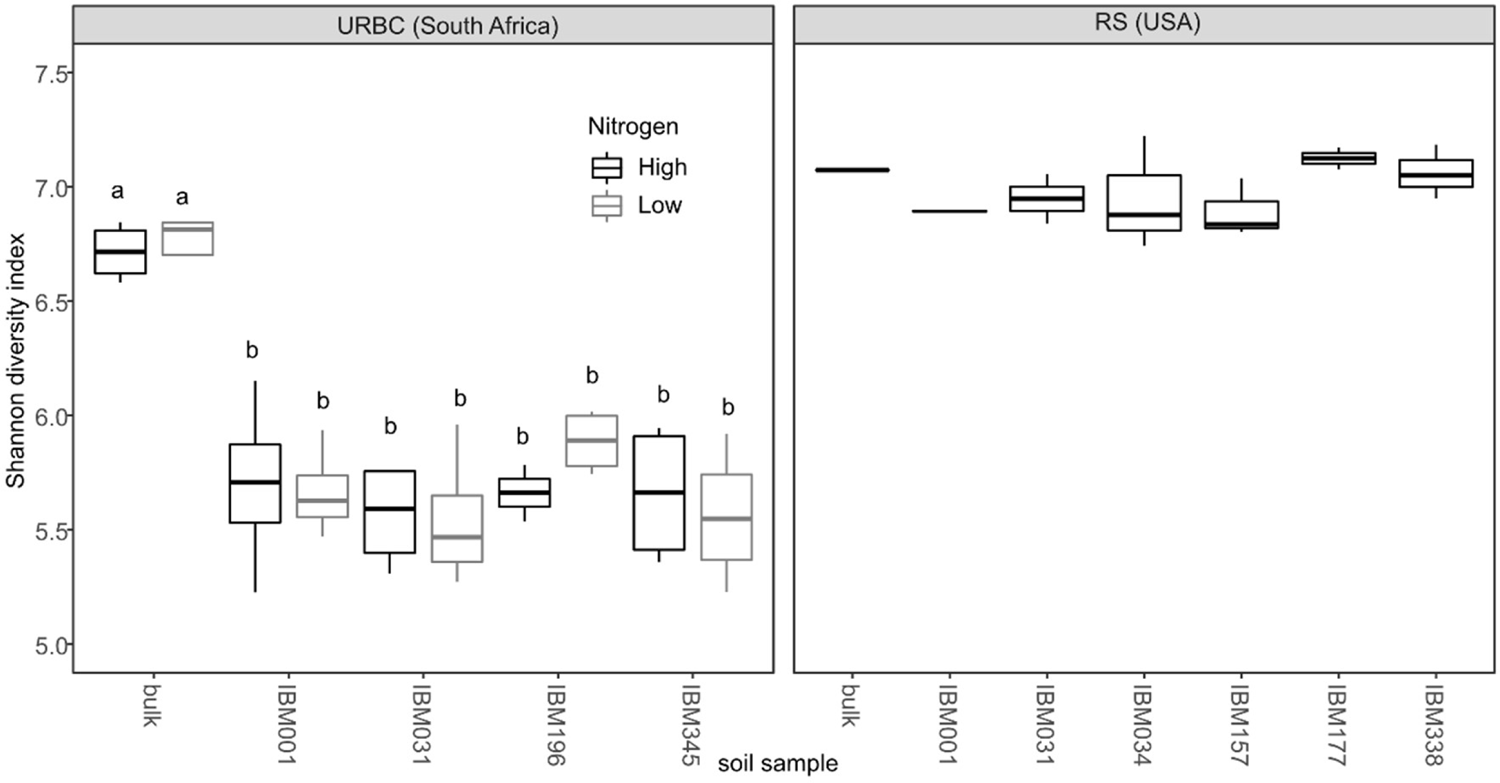
Alpha diversity of rhizosphere soil collected from all genotypes and bulk soil at the two sites. Horizontal box lines correspond to 25th, 50th, and 75th percentile; ranges are indicated by whiskers. For each boxplot n = 2-4. Boxes with the same letters indicate no significant differences in Shannon diversity indexes according to a LSD test with *P*<0.05. No significant differences were found between genotypes (and bulk soil) at RS.

### Nitrogen, genotype, RCA and rhizosphere effect on microbial communities

Microbial communities separated mainly by site (PCoA 1, Fig. 3) although there was a remarkable overlap of the community structure of bulk soil at URBC with bulk and rhizosphere soil communities at RS. Bulk soil samples ordinated apart from rhizosphere regardless of the nitrogen level at URBC (Supplementary Fig. 2). Genotype was not a significant grouping factor at either of the two locations (Fig. 3, Supplementary Fig. 2, Table 1). Transformed RCA values into qualitative ranks (Supplementary Table 3) had not significant effect on the weighted UniFrac distance at neither of the two sites (Table 1). RCA explained a greater amount of variation (9.5%) compared to genotype (7.6%) at URBC but the opposite was found at RS (Table 1). There was a significant effect of soil type (rhizosphere vs. bulk soil) at the two sites (Supplementary Table 2) but the percent of variation in beta diversity explained by soil type was over three times greater at URBC compared to RS. This stronger rhizosphere effect at URBC can be seen by comparing weighted-UniFrac distances between RCA phenotypes (separated by ranks) and bulk soils (Supplementary Fig. 3).

**Fig. 3.**
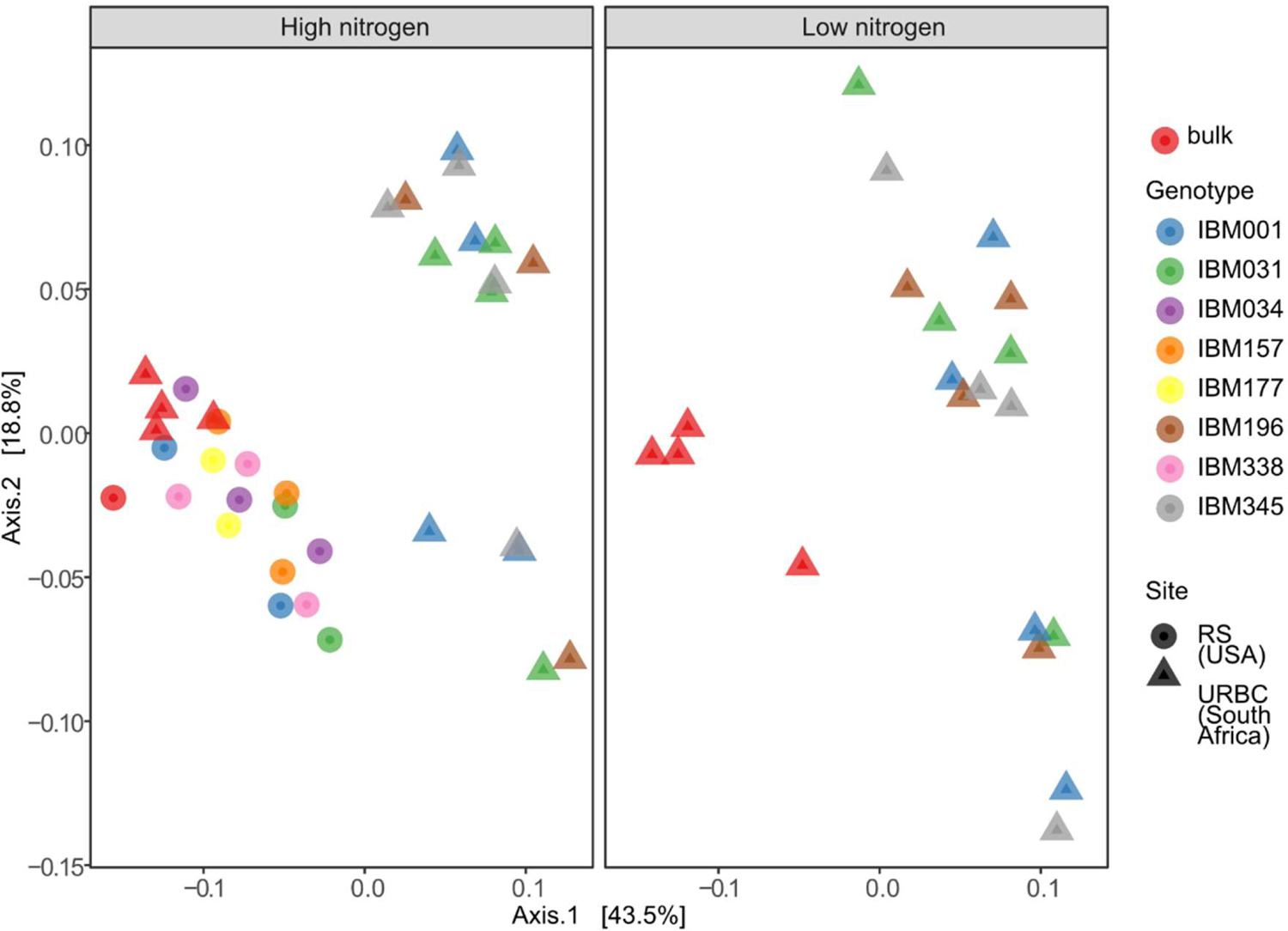
PCoAs using weighted UniFrac distances of the two sites, differentiated by genotypes. Nitrogen levels are in separate plots for URBC.

**Table 1.** Permutational MANOVA results using weighted UniFrac as a distance metric for the experiments by site. The model for URBC (South Africa) was weighted UniFrac distance ∼ Genotype*Nitrogen*RCA, and for RS (USA) the model was weighted UniFrac distance ∼ Genotype*RCA; with Block as random effect (with the option strata of the function *adonis* of the package *vegan* in R). For RCA (root cortical aerenchyma), we used qualitative ranks (See Supplementary Table 3 to see the values of each rank and Fig. 4 for data distribution within ranks). Bulk soil samples were excluded.

| Site | Factor | SS | % var<br>explained | R<sup>2</sup> | P<br>value |
| --- | --- | --- | --- | --- | --- |
| URBC (South Africa) | Genotype | 0.2775 | 7.6 | 0.07598 | 0.839 |
|  | Nitrogen | 0.1218 | 3.3 | 0.03335 | 0.055 |
|  | RCA | 0.3455 | 9.5 | 0.09459 | 0.401 |
|  | Genotype:Nitrogen | 0.2841 | 7.8 | 0.7778 | 0.785 |
|  | Genotype:RCA | 0.779 | 21.3 | 0.213 | 0.487 |
|  | Nitrogen:RCA | 0.295 | 8.1 | 0.0808 | 0.1 |
|  | Genotype:Nitrogen:RCA | 0.091 | 2.5 | 0.0249 | 0.804 |
|  | Residuals | 1.458 | 39.9 |  |  |
|  | Total | 3.6523 | 100.0 |  |  |
| RS (Pennsylvania) | Genotype | 0.477 | 30.042 | 0.30042 | 0.569 |
|  | RCA | 0.105 | 6.593 | 0.06593 | 0.426 |
|  | Genotype:RCA | 0.646 | 40.705 | 0.40705 | 0.319 |
|  | Residuals | 0.359 | 22.660 |  |  |
|  | Total | 1.586 | 100.000 |  |  |

### Effect of RCA on microbial communities

Contrasting RCA phenotypes were found among the plants evaluated (Fig. 4, Supplementary Table 3). Ordination plots of UniFrac distances by RCA (ranked according to Supplementary Table 3 and Fig. 4) show microbial communities of high and low RCA separated in the first component (Fig. 5). Ranges and other descriptive statistics of the complete set of anatomical and architectural phenes measured are provided in Supplementary Table 4 and Supplementary Table 5. All such phenes were used to select significant models and the resulting models are presented in Table 2. The effect of specific root phenes on rhizosphere microbial communities depended on site, and different models were selected with random permutations (Table 2). The selected models were then used with CCA (controlling for genotype and block - and nitrogen at URBC) to assess the effect of quantitative phene values on the unweighted and weighted UniFrac distances. Among the anatomical and architectural phenes measured at URBC (Table 2), RCA was significant for the unweighted (*P*=0.045, Table 2) and weighted (*P*=0.092, Table 2) UniFrac distances; and BottomAngle was significant (*P*=0.068, Table 2) for unweighted UniFrac distance, while D10 was significant (*P*=0.062, Table 2) for the weighted UniFrac distance. Among the anatomical variables measured at RS no significant models (with *P* < 0.1) were found for the weighted or the unweighted UniFrac distances. Diversity of rhizosphere microbial communities was greater in high RCA compared to low RCA under low nitrogen at URBC (Fig. 6). The effect of RCA on specific OTUs at the two experimental sites was further investigated.

**Fig. 4.**
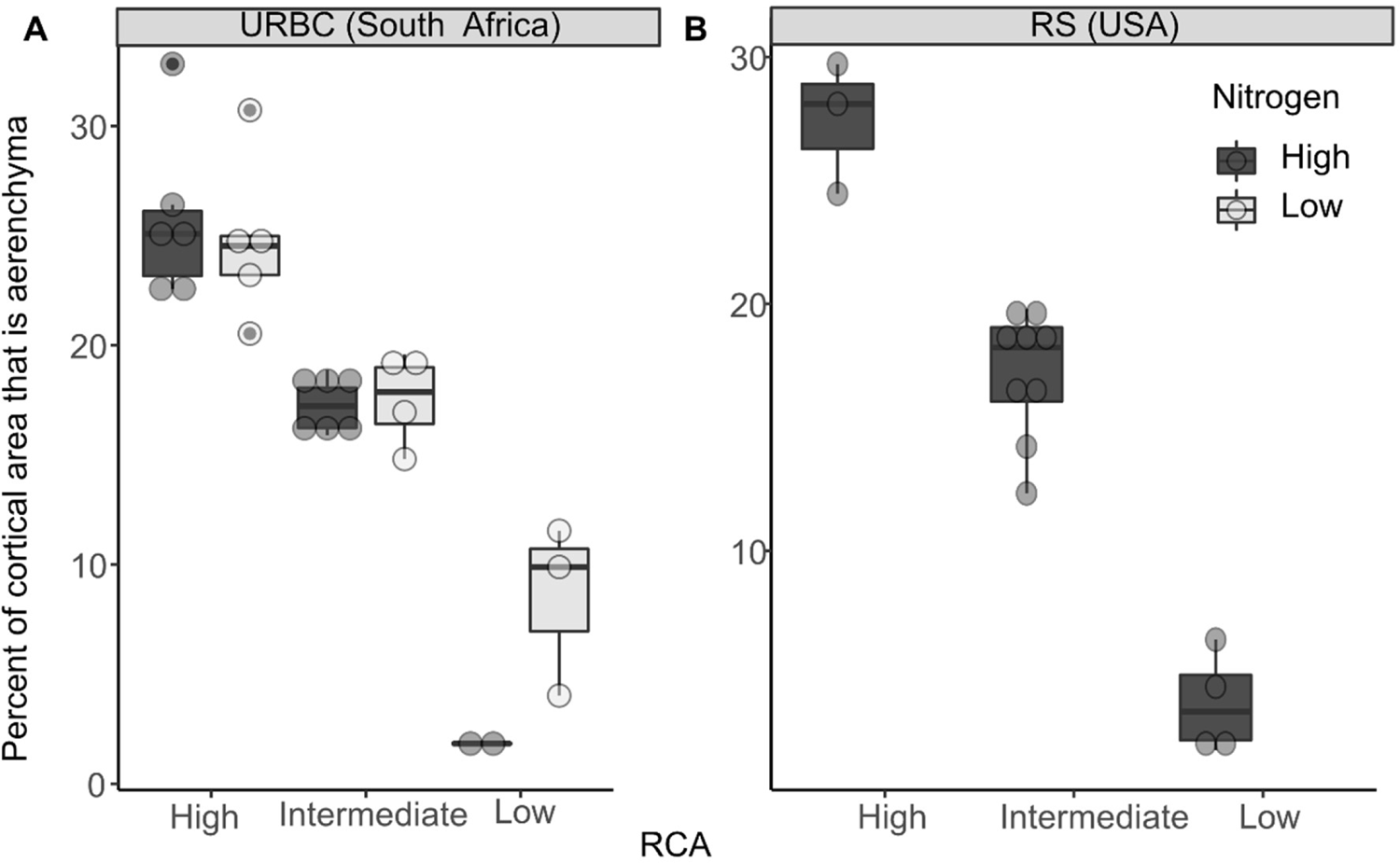
Bloxplots of percent cortical area that is aerenchyma (percCisA) by RCA ranks at the two sites. Horizontal box lines correspond to 25th, 50th, and 75th percentile; ranges are indicated by whiskers and points out of the boxes are outliers. For each boxplot the data points are indicated in open circles.

**Fig. 5.**
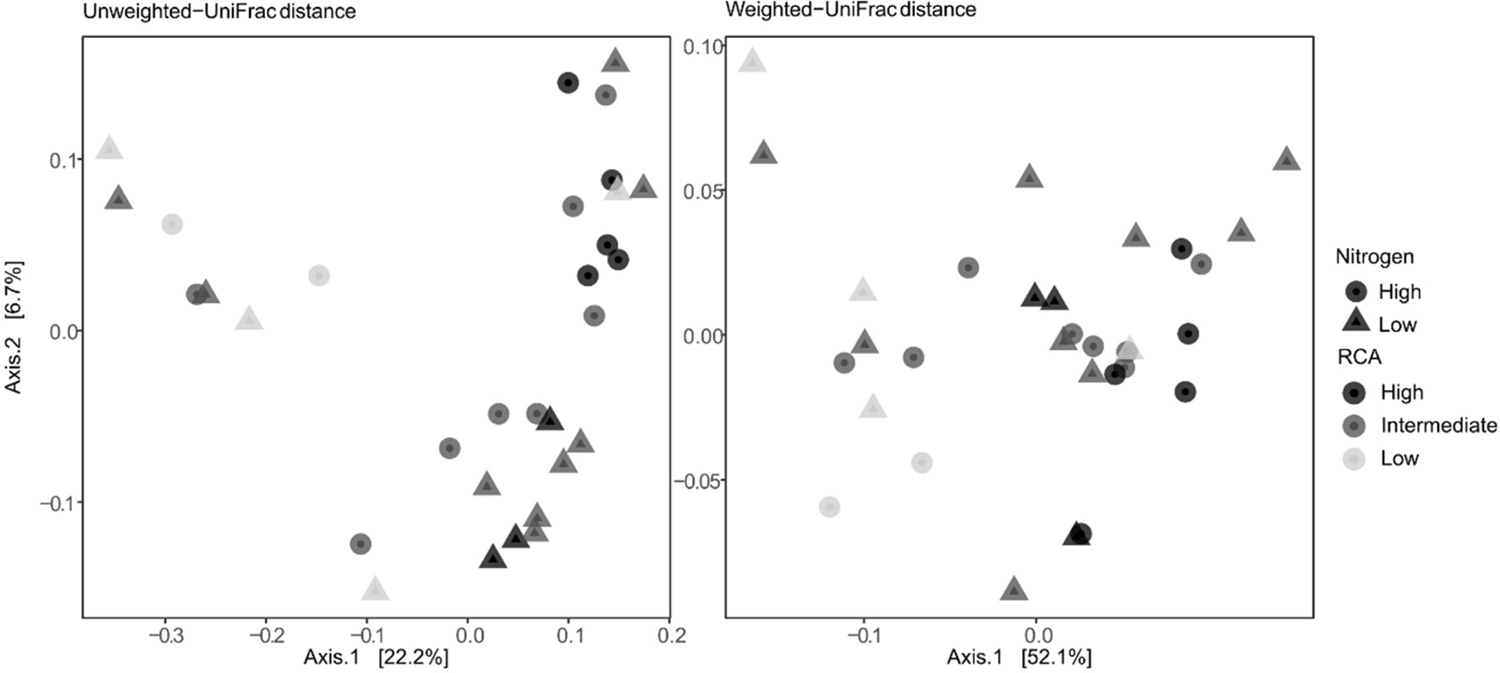
PCoAs using weighted and unweighted UniFrac distances by RCA qualitative ranks (determined as shown in Supplementary Table 3 and Fig.4), and additionally by nitrogen level at URBC.

**Fig. 6.**
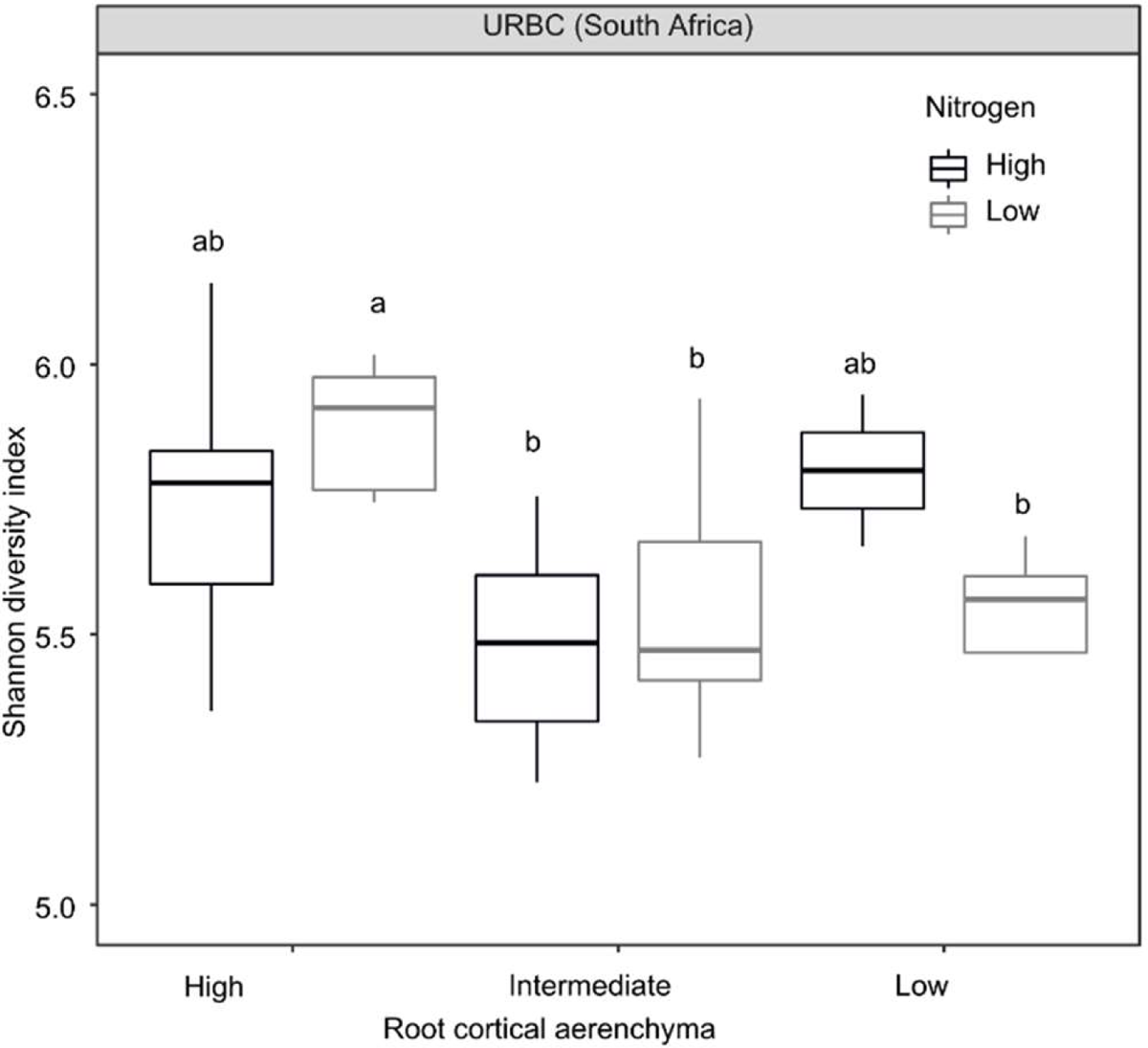
Boxplots of Shannon diversity values per RCA phenotype, and per nitrogen levels on rarefied data from URBC. Horizontal box lines correspond to 25th, 50th, and 75th percentile; ranges are indicated by whiskers. Letters indicate significant differences with a LSD test *P*<0.05.

**Table 2.**
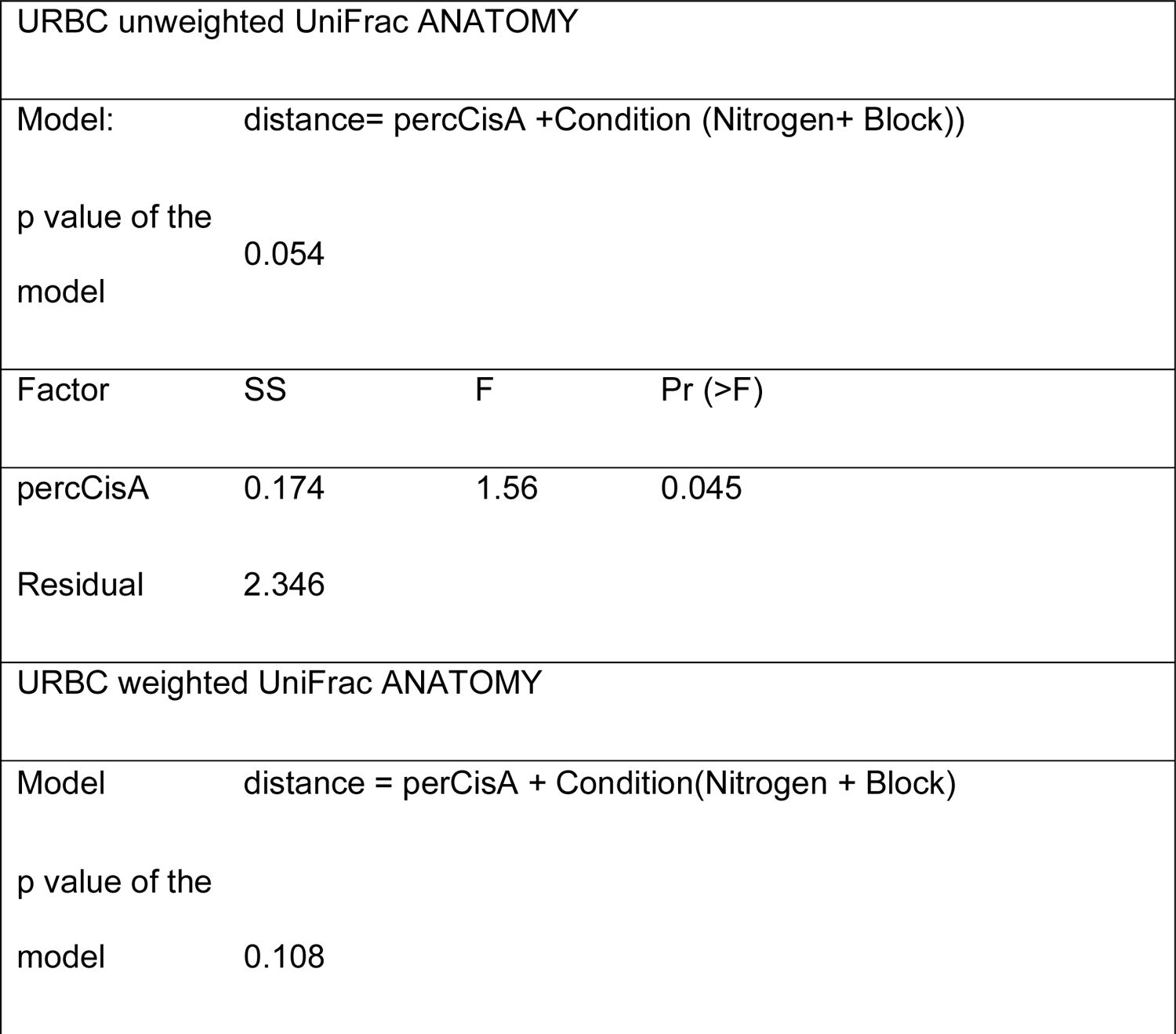
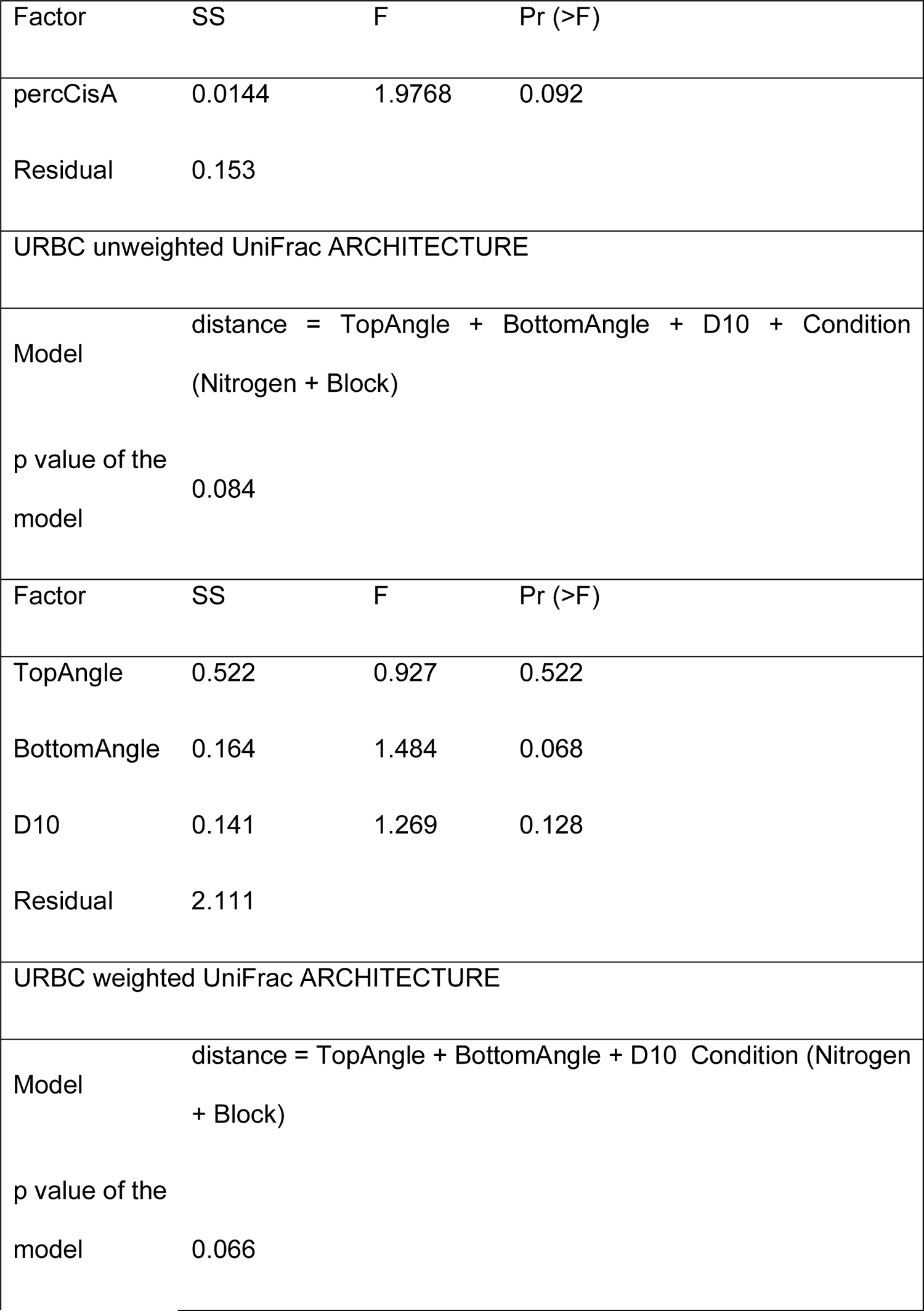
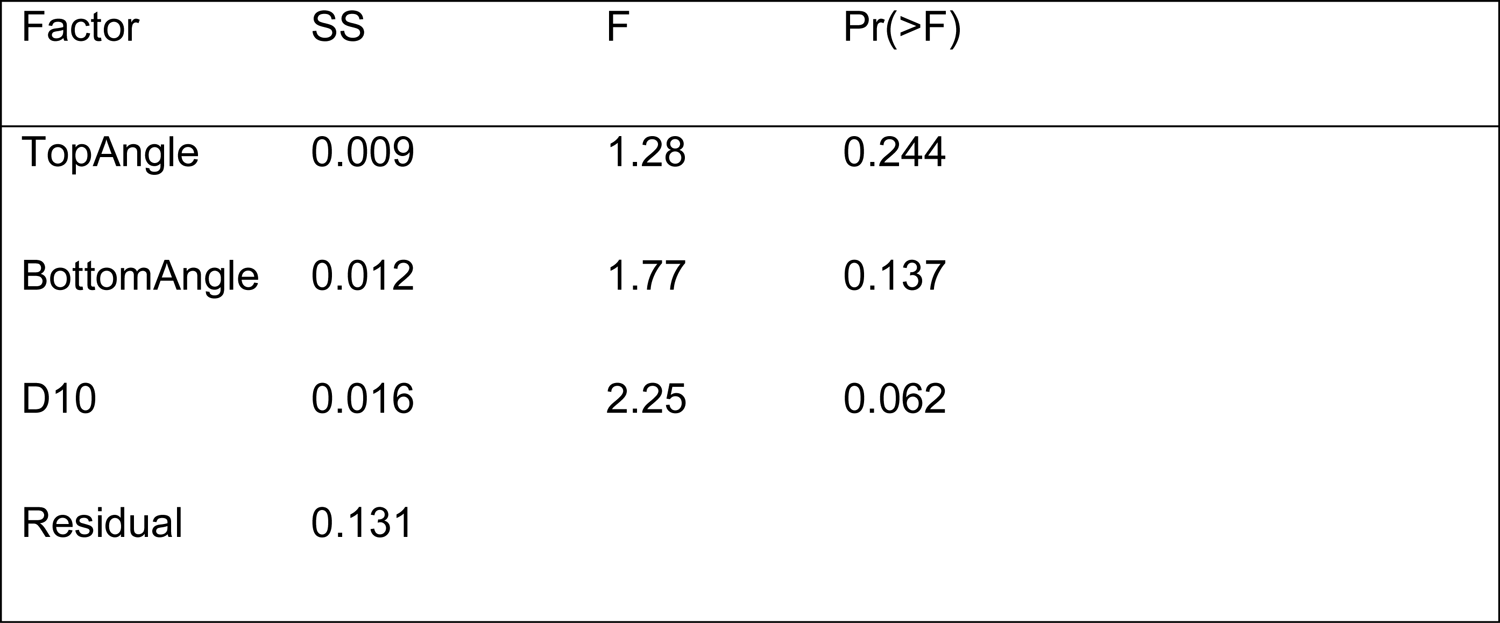
Models of unweighted and weighted UniFrac distances as functions of anatomical and architectural phenes at URBC (South Africa) and significance per phene, as selected by random permutations. Constrained correspondence analyses (CCA) were constrained by the factors Nitrogen, Genotype and Block. percCisA: Percentage of cortex that is aerenchyma. TopAngle: Angle along the outline of the root at 10% width accumulation. BottomAngle: Angle along the outline of the root at 70% width accumulation. D10: Accumulated width over the depth at 10% of the central path length.

| URBC unweighted UniFrac ANATOMY |  |  |  |
| --- | --- | --- | --- |
| Model: | distance= percCisA +Condition (Nitrogen+ Block)) |  |  |
| p value of the model | 0.054 |  |  |
| Factor | SS | F | Pr (>F) |
| percCisA | 0.174 | 1.56 | 0.045 |
| Residual | 2.346 |  |  |
| URBC weighted UniFrac ANATOMY |  |  |  |
| Model | distance = perCisA + Condition(Nitrogen + Block) |  |  |
| p value of the model | 0.108 |  |  |

| Factor | SS | F | Pr (>F) |
| --- | --- | --- | --- |
| percCisA | 0.0144 | 1.9768 | 0.092 |
| Residual | 0.153 |  |  |
| URBC unweighted UniFrac ARCHITECTURE |  |  |  |
| Model | distance = TopAngle + BottomAngle + D10 + Condition<br>(Nitrogen + Block) |  |  |
| p value of the<br>model | 0.084 |  |  |
| Factor | SS | F | Pr (>F) |
| TopAngle | 0.522 | 0.927 | 0.522 |
| BottomAngle | 0.164 | 1.484 | 0.068 |
| D10 | 0.141 | 1.269 | 0.128 |
| Residual | 2.111 |  |  |
| URBC weighted UniFrac ARCHITECTURE |  |  |  |
| Model | distance = TopAngle + BottomAngle + D10 Condition (Nitrogen<br>+ Block) |  |  |
| p value of the<br>model | 0.066 |  |  |

| Factor | SS | F | Pr(>F) |
| --- | --- | --- | --- |
| TopAngle | 0.009 | 1.28 | 0.244 |
| BottomAngle | 0.012 | 1.77 | 0.137 |
| D10 | 0.016 | 2.25 | 0.062 |
| Residual | 0.131 |  |  |

### Phenotype-sensitive OTUs

High-RCA plants had specific sets of rhizosphere prokaryotes that were significantly enriched or depleted compared to low-RCA plants (Fig. 7, Supplementary Fig. 4, Supplementary Fig. 5). At URBC, high-RCA rhizospheres hosted 95 significantly enriched OTUs and had 40 significantly decreased OTUs compared to low-RCA rhizospheres under high nitrogen. A similar ratio between enriched (43) and decreased (16) OTUs was found when comparing high and low RCA under low nitrogen (Fig. 7) at URBC. Also, alpha diversity between plant phenotypes (Fig. 6) show an increase of number of OTUs associated with high RCA phenotypes at URBC and low nitrogen, as well as the specific abundance values of OTUs by phylum and family (Supplementary Fig. 6 and Supplementary Fig. 7). When compared to bulk soil, rhizospheres of high-RCA plants had also a greater number of significantly enriched OTUs than low-RCA rhizospheres (Supplementary Fig. 4a), as well as a greater number of unique significantly enriched and decreased OTUs (Supplementary Fig. 4b). RCA phenotype had a weaker effect on the rhizosphere communities under high nitrogen at RS as there were fewer significantly enriched and depleted OTUs associated to each RCA phenotype (Fig. 7). The weaker rhizosphere effect at RS is also evident when the microbial communities of contrasting RCA phenotypes were compared to bulk soil (Supplementary Fig. 3 and Supplementary Fig. 5).

**Fig. 7.**
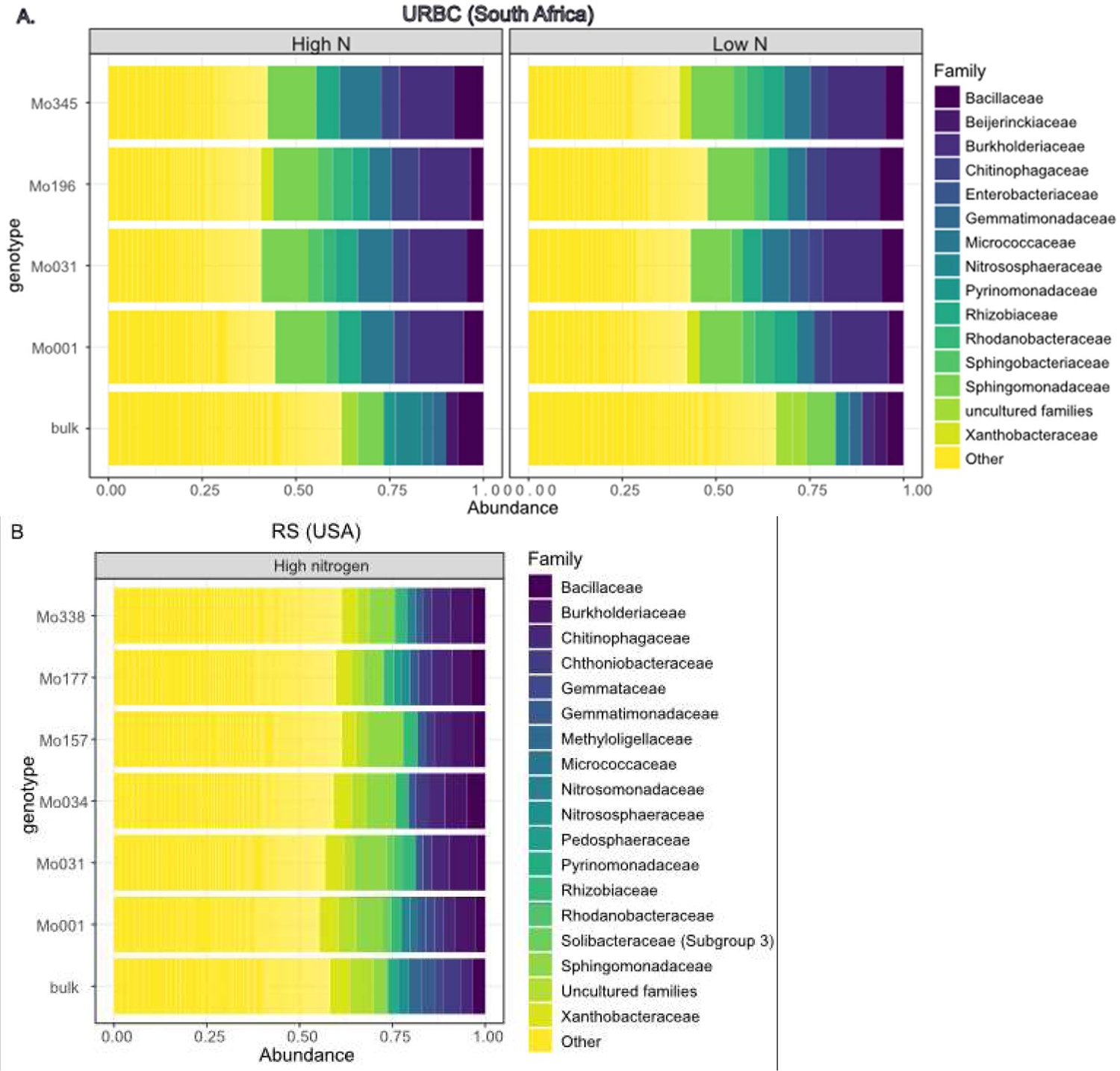
A: Abundance log change (y axis) of all the OTUs when RCA levels were compared within each RCA level. Black points indicate differentially enriched and depleted OTUs according to a likelihood ratio test with *P*<0.01, and grey points were non-differentially abundant between the two types of samples, Number of OTUs significantly enriched or decreased at each condition are in parenthesis. B: Number of the differentially enriched and depleted OTUs between each RCA level and nitrogen level at the two sites.

At URBC, RCA-sensitive OTUs from high RCA phenotypes shared some common features between high and low nitrogen. Significantly enriched microorganisms in high-RCA under high nitrogen (as shown in Fig. 7) belonged mainly to the phyla *Proteobacteria* (29.9% of the total enriched OTUs), *Acidobacteria* (27%), *Actinobacteria* (11.34%), *Bacteroidetes* (11.34%), and *Thaumarchaeota* (11.34%) and in minor proportions to the phyla *Chloroflexi* and *Firmicutes* (Supplementary Fig. 6a), similar to the distribution of the enriched OTUs in high-RCA rhizospheres of low nitrogen plots with *Proteobacteria* (34%), *Chloroflexi* (20%), *Bacteroidetes* (16%), and *Acidobacteria* (14%) and smaller proportions of *Actinobacteria* and *Firmicutes* (Supplementary Fig. 6b, Supplementary File S1). At the family level, High-RCA plants of the two nitrogen levels at URBC had some families significantly-enriched in common (although OTUs did not overlap) such as *Beijerinckiaceae, Burkholderiaceae, Haliangiaceae, Longimicrobiaceae, Microscillaceae, Chitinophagaceae, Bacillaceae, Paenibacillaceae, Polyangiaceae*, *Solibacteraceae*, and *Thermoanaerobaculaceae* (Supplementary Fig. 7).

There was greater diversity among the enriched OTUs of high-RCA plants at high nitrogen compared to low nitrogen at URBC. Enriched OTUs from the phyla *Thaumarchaeota*, *Gemmatimonadetes*, *Planctomycetes*, and *Dependentiae* in high-RCA plants were associated with high nitrogen, while an enrichment of OTUs from the phyla *Armatimonadetes* and *Cyanobacteria* were associated with low nitrogen (Supplementary Fig. 6). Several families were unique to high-RCA plants at each nitrogen level (Supplementary Fig. 7). Among the OTUs with the greatest abundance at high-RCA in high nitrogen, the families *Burkholderiaceae*, *Sphingomonadaceae*, *Nitrososphaeraceae*, had the most abundant significantly enriched OTUs compared to low-RCA rhizospheres (Supplementary Fig. 7). The most abundant OTUs enriched at low-RCA and high nitrogen belonged to the families *Burkholderiaceae* and *Chitinophagaceae*, and *Sphingobacteriaceae*. With low nitrogen and high-RCA, *Burkholderiaceae* had the most abundant OTUs. The families *Beijerinckiaceae*, *Bacillaceae*, *Burkholderiaceae* and *Chitinophagaceae* had OTUs that were shared between high and low nitrogen in high RCA plants, and that were among the most abundant and significantly enriched OTUs (Supplementary Fig. 7).

High-RCA rhizosphere of high-nitrogen plots at RS had 35 enriched OTUs. There was one shared OTU between URBC and RS that was depleted in high RCA roots (OTU 11202, an unclassified species of the family *Rhodanobacteraceae* (order *Xanthomonadales*, phylum Proteobacteria), (Fig. 7b). Enriched OTUs of high-RCA in RS were distributed among *Planctomycetes* (39% of the total enriched OTUs), *Acidobacteria* (17%) and *Proteobacteria* and *Actinobacteria* (14%), and in smaller proportions among *Firmicutes* (8%), and other phyla (<8%) (Supplementary Fig. 8, Supplementary File S1); whereas the 45 enriched OTUs of low-RCA rhizospheres at RS had a contrasting phyla distribution with *Proteobacteria* (44%), *Bacteroidetes* (16%) as dominant phyla followed by *Actinobacteria* (13%), *Planctomycetes* (9%) and *Acidobacteria* (6%) and other phyla (<6%). The most abundant OTUs in RS belonged to the genus *Oceanobacillus* (family *Bacillaceae*) in high-RCA plants, and to the genus *Aquicella* (family *Diplorickettsiaceae*) in low-RCA plants.

## DISCUSSION

Contrasting levels of RCA of field-grown maize were associated with specific compositions of the rhizosphere microbiome and the extent of this association depended on the geographic location, being more defined at URBC (South Africa). Regardless of the nitrogen fertilization treatment, RCA was associated with a set of significantly enriched OTUs under an intensively managed agricultural sandy soil in South Africa but had no significant effect on the rhizosphere microbial richness in a finer-textured soil of Pennsylvania (USA). Our results indicate that root phenotypes may explain part of the variability in the rhizosphere microbial composition and constitute a starting point to further study root phenotype effects on the root microbiome of agricultural species.

The dominant phyla *Proteobacteria*, *Acidobacteria*, *Bacteroidetes* and *Actinobacteria* found here overlap those reported among the dominant phyla in agricultural and rhizosphere soils (Lundberg et al. 2012; Peiffer et al. 2013; Philippot et al. 2013; Sul et al. 2013). The stronger rhizosphere effect observed at URBC (South Africa) compared to RS (USA) could be explained by differences in soil properties and agricultural management of the two sites. The sandy, low organic matter soil at URBC may have offered a more restrictive environment for microbial growth compared to the more fertile silt-loam at RS (Supplementary Table 1). Additionally, crop rotations at RS in comparison with maize after maize monoculture at URBC could have provided more diverse microbial communities at RS (Fig. 2). Therefore, root phenotypes had a significant impact on soil microorganisms at URBC (Supplementary Fig. 3, Supplementary Fig. 4), where soils contain less organic matter (<0.5%, Supplementary Table 1) and are coarser in texture than RS, leading to less well-developed soil structure. Better soil structure (e.g. more aggregates) creates more diverse microenvironments and contributes to greater microbial diversity (Fierer 2017; Sexstone et al. 1985). Likewise, greater organic matter content can be associated with greater microbial diversity due to the more diverse carbon sources for microbial decomposition (Sul et al. 2013). Accordingly, RCA separated the microbial communities at URBC, where there was a significant rhizosphere effect (Fig. 5, Supplementary Fig. 3, and Supplementary Table 2), while no separation by RCA levels was observed at RS. Despite the differences in rhizosphere effects between the two sites, it is noteworthy to find two enriched (*Proteobacteria* and *Bacteroidetes*) and two depleted (*Acidobacteria* and *Nitrospirae*) phyla in maize rhizospheres of the two sites, in accord with previous studies in which *Proteobacteria* and *Bacteroidetes* were enriched (Bakker et al. 2015; Peiffer et al. 2013) and *Acidobacteria* depleted in the rhizosphere soil (Fierer et al. 2007; Niu et al. 2017; Peiffer et al. 2013). Among the significantly depleted or enriched OTUs at contrasting levels of RCA found here, the genera *Agromyces*, *Bacillus*, *Caulobacter*, *Chthoniobacter*, *Flavobacterium*, *Nocardioides* and *Sphingomonas* were recently reported as part of the maize core microbiome (Walters et al. 2018). These findings demonstrate the intrinsic selectivity of soil microbial communities and the potential importance of RCA (and possibly other root phenes) on changing the composition of the communities in the rhizosphere. Additionally, and similar to our results, Dohrmann et al. (2013) found that nitrifiers were slightly enriched in the rhizosphere of genetically modified Bt field-grown maize but they found bacteria (*Nitrosomonas* and *Nitrospira*) as opposed to the ammonia oxidizing archaeans of the family *Nitrososphaereaceae* found here. Dohrmann et al. (2013) suggested that their enrichment of nitrifying bacteria could be linked to a greater protein content in Bt maize due to the possible overexpression of Cry proteins. Since ammonia oxidizing archaeans outcompete ammonia oxidizing bacteria under lower ammonia concentration (Hatzenpichler 2012), our results at URBC (sandy, low organic matter content and low pH soil in South Africa) may correspond to rhizospheres with low nitrogen concentration in high-RCA plants, even under high nitrogen fertilization. Moreover, enrichment of archaea of the *Nitrososphaeraceae* family is highly suggestive of changes in nitrification as members of this family are obligately aerobic chemolithoautotrophs capable of nitrification. They can be mixotrophic, requiring organic substrates for growth, and this may contribute to their enrichment in high RCA plants. The mechanisms underlying this merit further investigation.

Our results support the hypothesis that root control of rhizosphere communities among related genotypes is associated with root phenotypes (Fig. 5, Table 1, Table 2, Fig. 6). We propose that phenotypes have a stronger, and perhaps more predictive effect on microbial communities compared to genotype effects as observed at URBC, with genotypes explaining 7% of variation and RCA explaining 9% of the variation (Table 1), this not accounting for other phenes that, in addition to RCA, might have significant effect on rhizosphere biodiversity. This hypothesis will need additional support. However, it is noteworthy that genotypic effect of related maize lines has shown modest or partial effects on rhizosphere microbial communities in the present and in previous studies (Bakker et al. 2015; Dohrmann et al. 2013; Fang et al. 2005; Peiffer et al. 2013; Walters et al. 2018), and no exploration of root phenotypes has been reported to our knowledge in large-scale microbiome studies of rhizospheres. Moreover, there appear to be root phenes that are more important than others as drivers of the microbial composition in the rhizosphere as shown by the CCA analysis - when controlling for nitrogen and genotype effects (Table 2). The root phenes shaping the rhizosphere communities varied by site, with RCA significant for the UniFrac distance metrics at URBC. While the differences in RCA at RS had no significant effects on rhizosphere microbial diversity, we found a few uniquely enriched taxa with each phenotype (high and low-RCA) (Fig. 7), with the most abundant taxa (genus *Oceanobacillus*) being associated with plant growth promotion (Supplementary File S1, Supplementary results).

We propose that the diversity associated with contrasting levels of RCA observed at URBC may be associated with changes in functions in the rhizosphere microbial community. More specifically, two possible mechanisms that could be further studied as factors affecting rhizosphere microbial diversity are the diffusion of oxygen from aerenchyma lacunae and the rhizodeposition of carbon as influenced by RCA. Greater oxygen concentration and possibly, differences in carbon rhizodeposition into the rhizosphere of the high-RCA plants may be associated with the ∼2 fold greater number of significantly enriched OTUs observed in high-RCA plants phenotypes compared to low-RCA plants at URBC under high and low nitrogen (Fig. 7). Plants with increased RCA may have reduced carbon rhizodeposition in axial roots as a consequence of the loss of cortical tissue; this effect may be intensified under low nitrogen given the overall low nitrogen content of the plant. However, it is also possible that under low nitrogen, plants with increased RCA have more carbon to invest in rhizodeposition compared to reduced-RCA plants as an indirect consequence of the benefits of RCA on nitrogen acquisition under low nitrogen (Saengwilai et al. 2014). Reduced cortical tissue of high-RCA plants reduces the metabolic burden of soil exploration and nutrient capture (Lynch 2015).

RCA had significant effects on the abundances the ammonia oxidizing archaean family *Nitrososphaeraceae* in high-RCA plants growing under high nitrogen at URBC (Supplemental File S1). Enrichment of the archaean *amoA* genes (genes encoding for the ammonia monooxygenase) were previously found in a study with field-grown maize rhizosphere (Li et al. 2014) in accordance with our findings. High abundances of *Nitrososphaeraceae* could cause a net decrease of ammonia in the rhizosphere, forcing other microbial species or even the plant itself to invest reductive power in nitrate assimilation or could also promote nitrogen losses from the rhizosphere if the nitrate generated is lost as leachate or ultimately converted into gaseous nitrogen (Stahl and Torre 2012). The enrichment of *Nitrososphaeraceae* in high-RCA and high-N plants at URBC could indicate a low continuous supply of ammonium in accordance with previous research indicating that archaeal ammonia oxidizers are adapted to lower ammonia availability (Hatzenpichler 2012; Stahl and Torre 2012; Sterngren et al. 2015). However, since our analyses are based on taxonomy these hypotheses merit further research.

The present study provides insights into the effects of RCA on rhizosphere microbial communities of maize grown in two contrasting environments, and give rise to interesting questions and hypotheses for future research. Further research could be conducted to reveal more detailed effects exploring more root phenotypes and expanding to root architectural phenes. Here, we found that together with RCA, rooting angle had significant effects on the communities in the maize rhizosphere at URBC (Table 2). The use of larger sets of genotypes as well as the study of the effects of phenotypes in combination with plant developmental stages will also add to our findings. Additionally, the inclusion of eukaryotes is crucial for the understanding of the effects of aerenchyma and other phenotypes on the fungal populations closely related to the root cortex.

The selection of plants targeting root ideotypes that improve soil exploration under low-nutrient and drought stress would be benefited by a concomitant selection of beneficial microbiomes. This study is a pioneer in this endeavor by suggesting possible habitat changes provided by contrasting RCA and the associated microorganisms. Plant and microbiome breeding together could produce ideal combinations of roots and microbes adapted to resource scarcity to improve plant growth and productivity.

## ACKNOWLEDGEMENTS

Robert Snyder, Michael Williams, Gustavo da Silveira, Andrew Evensen, Melda Manchidi Shaku, Tsitso Zechariah Mokoena, Vincent Nkhumeleni Rambau, Javier Ceja Navarro, and Shi Wang provided technical assistance. The NVIDIA Corporation donated the Tesla K40 GPU used for this research. We thank Drs. Mary Ann Bruns and Alexander Bucksch for their discussion and comments.

## Statement of author contributions

J.P.L. and T.G.C. conceived and designed the research; T.G.C. conducted the experiments and performed statistical analysis; E.L.B. provided the facility to conduct the sequencing of DNA and provided technical and scientific assistance for the analysis and data interpretation; U.K. performed the UPARSE work of the raw sequences; C.R. performed the work with Qiime and provided technical assistance for the statistical analysis; T.G.C. and J.P.L. wrote the article with contributions of all the authors. J.P.L. agrees to serve as the author responsible for contact and ensures communication.

## Supplementary Tables

**Supplementary Table 1.** Summary of soil analyses performed at URBC (South Africa) and RS (USA) as an external service. Extraction methods: P - Bray I \ Olsen (pH >= 7.3), Cations – NH_4_OAc, Organic C - Walkley-Black method, Fe,Mn,Zn,Cu,Ni – DTPA, Tot-N - 0.1N K_2_SO_4_. NA: not available information.

| Measurement | Units | Values |  |  |
| --- | --- | --- | --- | --- |
|  |  | URBC |  | RS |
|  |  | LN | HN | HN |
| Bulk density | $\text{g}\cdot\text{cm}^3$ | 1.250-1.511 | | 1.400-1.600 |
| pH | KCl | 3.8-5.3 |  | 6.7-7.3 |
| S | $\text{mg}\cdot\text{kg}^{-1}$ | 7.0 - 89.0 | | 7.6-16.4 |
| P | $\text{mg}\cdot\text{kg}^{-1}$ | 16-73 | | 35-111 |
| K | $\text{mg}\cdot\text{kg}^{-1}$ | 46-70 | | 87-271 |
| Ca | $\text{mg}\cdot\text{kg}^{-1}$ | 86-240 | | 1165-2907 |
| Mg | $\text{mg}\cdot\text{kg}^{-1}$ | 26-57 | | 98-168 |
| ECEC | Calculated | 1.1-1.9 |  | 8.3-16.7 |
| Total N | $\text{mg}\cdot\text{kg}^{-1}$ | 6.0-13 | NA | NA |
| $\text{NO}_3$ | $\text{mg}\cdot\text{kg}^{-1}$ | 5.0-12 | NA | NA |
| $\text{NH}_4$ | $\text{mg}\cdot\text{kg}^{-1}$ | 1.0-3 | NA | NA |
| Organic matter content | % | 0.1-0.5 |  | 0.9-2 |

**Supplementary Table 2.** Permutational MANOVA results using weighted UniFrac as a distance metric for the experiments by site. The Adonis model for each experiment was: at URBC, weighted UniFrac distance ∼ Soil type * Nitrogen + Block, at RS weighted UniFrac distance ∼ Soil type + Block. Soil type refers to rhizosphere soil and bulk soil.

| Site | Factor | SS | % Explained | P value |
| --- | --- | --- | --- | --- |
| URBC (South Africa) | Soil type | 0.185 | 38.6 | 0.001 |
|  | Nitrogen | 0.010 | 2.1 | 0.224 |
|  | Block | 0.040 | 8.4 | 0.002 |
|  | Soil type:Nitrogen | 0.005 | 1.0 | 0.58 |
|  | Residuals | 0.239 | 49.9 |  |
|  | Total | 0.479 |  |  |
| RS (USA) | Soil type | 0.012 | 11.4 | 0.001 |
|  | Block | 0.031 | 29.5 | 0.001 |
|  | Residuals | 0.063 | 60.0 |  |
|  | Total | 0.105 |  |  |

**Supplementary Table 3.** Values (in percentage) of RCA expressed as percent of the cortical area that is aerenchyma (percCisA) measured at RS and URBC.

| Site | Phene states | Quantitative values |
| --- | --- | --- |
| URBC | High | >20 |
| (South Africa) | Intermediate | 12 - 20 |
|  | Low | <12 |
| RS (USA) | High | >20 |
|  | Intermediate | 10 - 20 |
|  | Low | <10 |

**Supplementary Table 4.** Descriptive statistics of anatomical and architectural phenes measured at URBC.

| Phene abbreviation | Description | Units | Median | Mean | min | max | Variance | Standard Deviation | Coefficient of Variation |
| --- | --- | --- | --- | --- | --- | --- | --- | --- | --- |
| RXSA | Root cross section area | mm<sup>2</sup> | 1.908 | 1.941 | 0.642 | 5.106 | 0.603 | 0.777 | 0.40 |
| TCA | Total cortex area | mm<sup>2</sup> | 1.423 | 1.426 | 0.454 | 3.657 | 0.310 | 0.557 | 0.39 |
| TSA | Total stele area | mm<sup>2</sup> | 0.495 | 0.537 | 0.172 | 1.644 | 0.066 | 0.257 | 0.48 |
| AA | Aerenchyma area | mm<sup>2</sup> | 0.240 | 0.248 | 0.000 | 0.597 | 0.022 | 0.149 | 0.60 |
| MXVA | Total metaxylem vessel area | mm<sup>2</sup> | 0.074 | 0.074 | 0.025 | 0.136 | 0.000 | 0.022 | 0.30 |
| MXVS | Mean metaxylem vessel size | mm<sup>2</sup> | 0.006 | 0.006 | 0.002 | 0.013 | 0.000 | 0.002 | 0.26 |
| MXVN | Metaxylem vessel number | count | 12.000 | 12.391 | 7.000 | 21.000 | 9.364 | 3.060 | 0.25 |
| CCFN | Cortical cell file number | count | 10.333 | 10.733 | 6.000 | 18.000 | 4.966 | 2.229 | 0.21 |
| LCA | Living cortical area | mm<sup>2</sup> | 0.221 | 0.263 | 0.107 | 1.093 | 0.028 | 0.168 | 0.64 |
| CCS | Cortical cell size | mm<sup>2</sup> | 0.000237 | 0.000259 | 0.000139 | 0.000456 | 0.000000 | 0.000082 | 0.315817 |
| C:S | Cortex:Stele ratio | dimensionless | 2.673 | 2.805 | 1.661 | 4.928 | 0.431 | 0.656 | 0.23 |
| perCisA | Percentage of cortex that is aerenchyma | % | 19.965 | 17.823 | 0.000 | 34.237 | 76.128 | 8.725 | 0.49 |
| perCisCC | Percentage of cortex that is LCA | % | 18.804 | 19.465 | 10.209 | 29.895 | 19.840 | 4.454 | 0.23 |
| perXSisCC | Percentage of cross section that is LCA | % | 13.977 | 14.538 | 7.429 | 25.025 | 13.362 | 3.655 | 0.25 |
| S_Diam | Stem diameter | mm | 14.852 | 14.376 | 6.988 | 18.771 | 6.020 | 2.454 | 0.17 |
| RootArea | number of pixels that belong to the root system | number of pixels | 16716.534 | 16598.836 | 7441.681 | 26802.388 | 21008886.169 | 4583.545 | 0.28 |
| RootDensity | Ratio between root and background pixels | root pixels*back ground pixels<sup>-1</sup> | 3.696 | 3.879 | 1.262 | 7.287 | 1.747 | 1.322 | 0.34 |
| TopAngle | Angle along the OTUline of the root at 10% width accumulation | Sexagesimal degree (°) | 7.289 | 22.820 | 0.061 | 68.237 | 583.971 | 24.165 | 1.06 |
| BottomAngle | Angle along the OTUline of the root at 70% width accumulation | Sexagesimal degree (°) | 26.228 | 25.418 | 0.552 | 44.612 | 134.372 | 11.592 | 0.46 |
| RootPaths | Number of paths detected for the root system. Correlated with number of root tips | Count | 542.000 | 594.031 | 249.000 | 1336.000 | 45409.843 | 213.096 | 0.36 |
| D10 | Accumulated width over the depth at 10% of the central path length. Closely related to the root-top angle for maize. | % | 0.340 | 0.334 | 0.173 | 0.453 | 0.004 | 0.066 | 0.20 |
| LatRootLength | Average length of the detected lateral roots emerging from the central path long of the excised roots | cm | 193.399 | 192.664 | 137.866 | 268.145 | 1057.415 | 32.518 | 0.17 |
| NodalRootLength | Length of the central path along the excised root | cm | 299.660 | 293.717 | 186.522 | 392.363 | 1999.905 | 44.720 | 0.15 |
| LatBD | Lateral branching density | lateral roots*cm<sup>-1</sup> | 11.985 | 12.436 | 1.293 | 31.368 | 56.974 | 7.548 | 0.61 |
| NodalRootDiam | Average nodal root diameter | µm | 174.402 | 170.041 | 13.120 | 294.623 | 3560.569 | 59.670 | 0.35 |
| DistFirstLat | Distance to first lateral | cm | 0.575 | 1.415 | 0.000 | 10.269 | 4.108 | 2.027 | 1.43 |

**Supplementary Table 5.** Descriptive statistics of anatomical phenes measured at RS.

| Phene abbreviation | Description | Units | min | max | Median | Mean | Standard Error | Variance | Standard Deviation | Coefficient of Variation |
| --- | --- | --- | --- | --- | --- | --- | --- | --- | --- | --- |
| RXSA | Root cross section area | mm<sup>2</sup> | 0.576 | 2.750 | 1.229 | 1.261 | 0.096 | 0.222 | 0.472 | 0.37 |
| TCA | Total cortex area | mm<sup>2</sup> | 0.400 | 2.149 | 0.956 | 0.959 | 0.077 | 0.142 | 0.377 | 0.39 |
| TSA | Total stele area | mm<sup>2</sup> | 0.158 | 0.601 | 0.273 | 0.302 | 0.022 | 0.011 | 0.107 | 0.35 |
| C:S | Cortex:Stele ratio | dimensionless | 1.816 | 4.951 | 3.316 | 3.200 | 0.142 | 0.481 | 0.693 | 0.22 |
| AA | Aerenchyma area | mm<sup>2</sup> | 0.017 | 0.522 | 0.129 | 0.151 | 0.022 | 0.012 | 0.110 | 0.72 |
| perCisA | Percentage of cortex that is aerenchyma | % | 1.930 | 29.710 | 16.498 | 15.645 | 1.562 | 58.559 | 7.652 | 0.49 |
| MXVA | Total metaxylem vessel area | mm<sup>2</sup> | 0.034 | 0.136 | 0.059 | 0.066 | 0.005 | 0.001 | 0.025 | 0.37 |
| MXVS | Mean metaxylem vessel size | mm<sup>2</sup> | 0.002 | 0.009 | 0.006 | 0.006 | 0.000 | 0.000 | 0.002 | 0.29 |
| MXVN | Metaxylem vessel number | count | 8 | 20 | 10.5 | 11.639 | 0.773 | 14.347 | 3.788 | 0.33 |
| CCFN | Cortical cell file number | count | 7 | 13 | 9.5 | 9.597 | 0.279 | 1.865 | 1.365 | 0.14 |
| LCA | Living cortical area | mm<sup>2</sup> | 0.132 | 0.579 | 0.351 | 0.347 | 0.027 | 0.017 | 0.131 | 0.38 |
| perCisCC | Percentage of cortex that is LCA | % | 26.943 | 49.813 | 36.288 | 36.646 | 1.413 | 47.899 | 6.921 | 0.19 |
| perXSisCC | Percentage of cross section that is LCA | % | 19.322 | 39.014 | 26.738 | 27.666 | 1.098 | 28.914 | 5.377 | 0.19 |
| CCS | Cortical cell size | mm<sup>2</sup> | 71.225 | 240.201 | 123.916 | 137.241 | 9.565 | 2195.808 | 46.859 | 0.34 |

## Supplementary Figures

**Supplementary Fig. 1.**
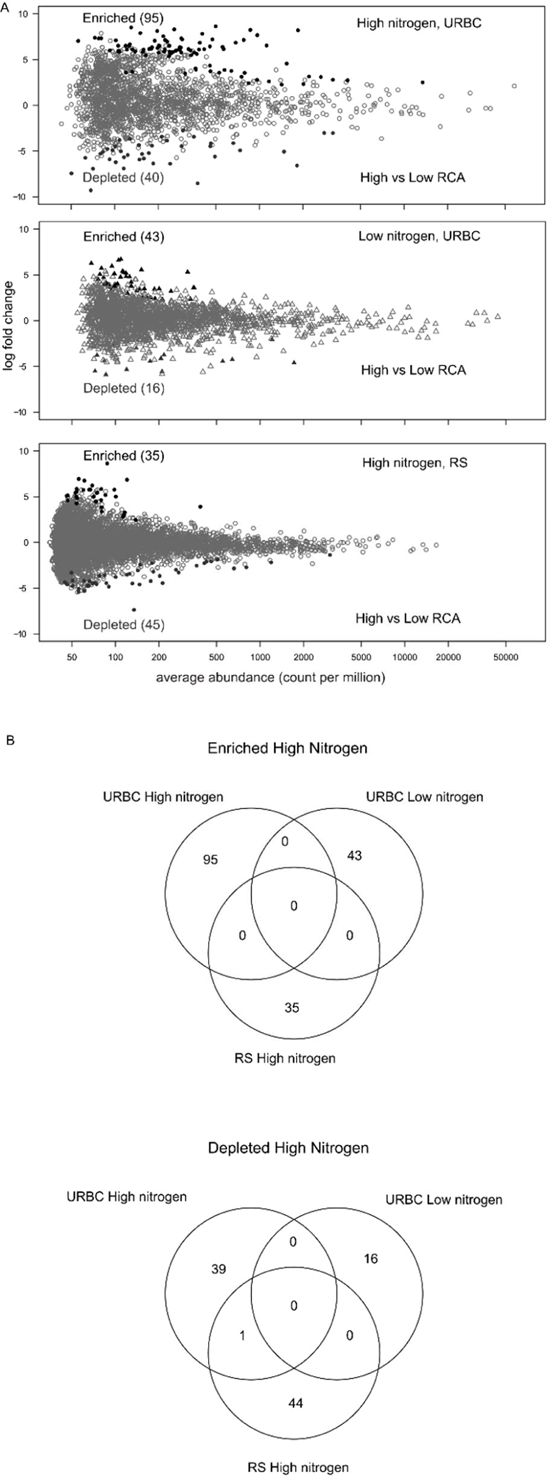
Bar plots of the relative abundances by family in each sample and experimental site. Families with abundances values below 0.03 (at URBC) and 0.02 (at RS) were grouped as “Other”. At RS (USA), uncultured families were distributed among 19 different phyla, among which *Actinobacteriaceae* and *Verrucomicrobia* were the most abundant. At URBC (South Africa), uncultured families were distributed among 14 phyla with *Actinobacteria* and *Acidobacteria* having the greatest abundance values.

**Supplementary Fig. 2.**
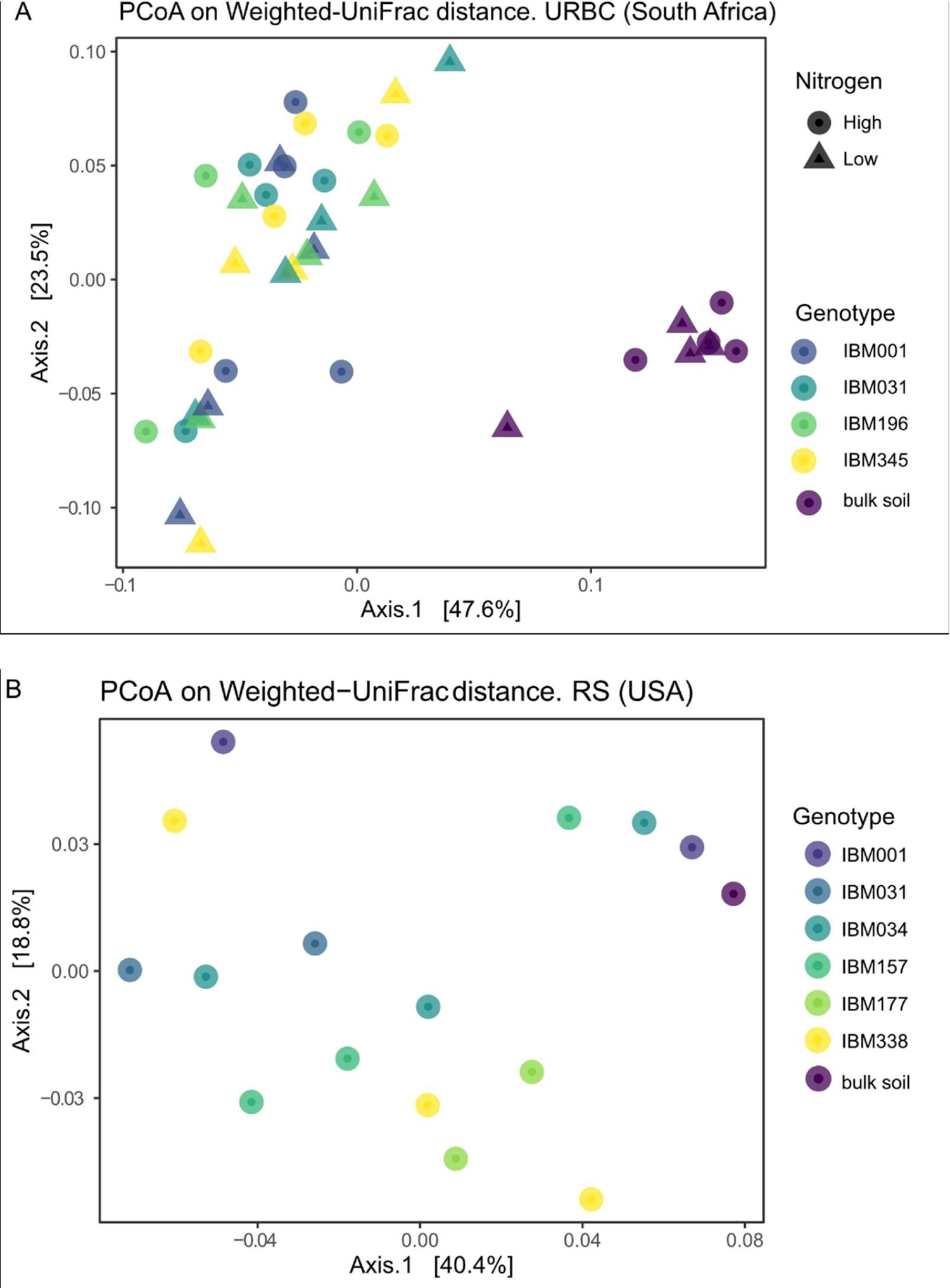
PCoAs using weighted UniFrac distances of each experimental site (A and B) by type of soil sample (rhizosphere vs bulk soil) in both sites, and additionally by nitrogen at URBC. Plots show the first two principal axes.

**Supplementary Fig. 3.**
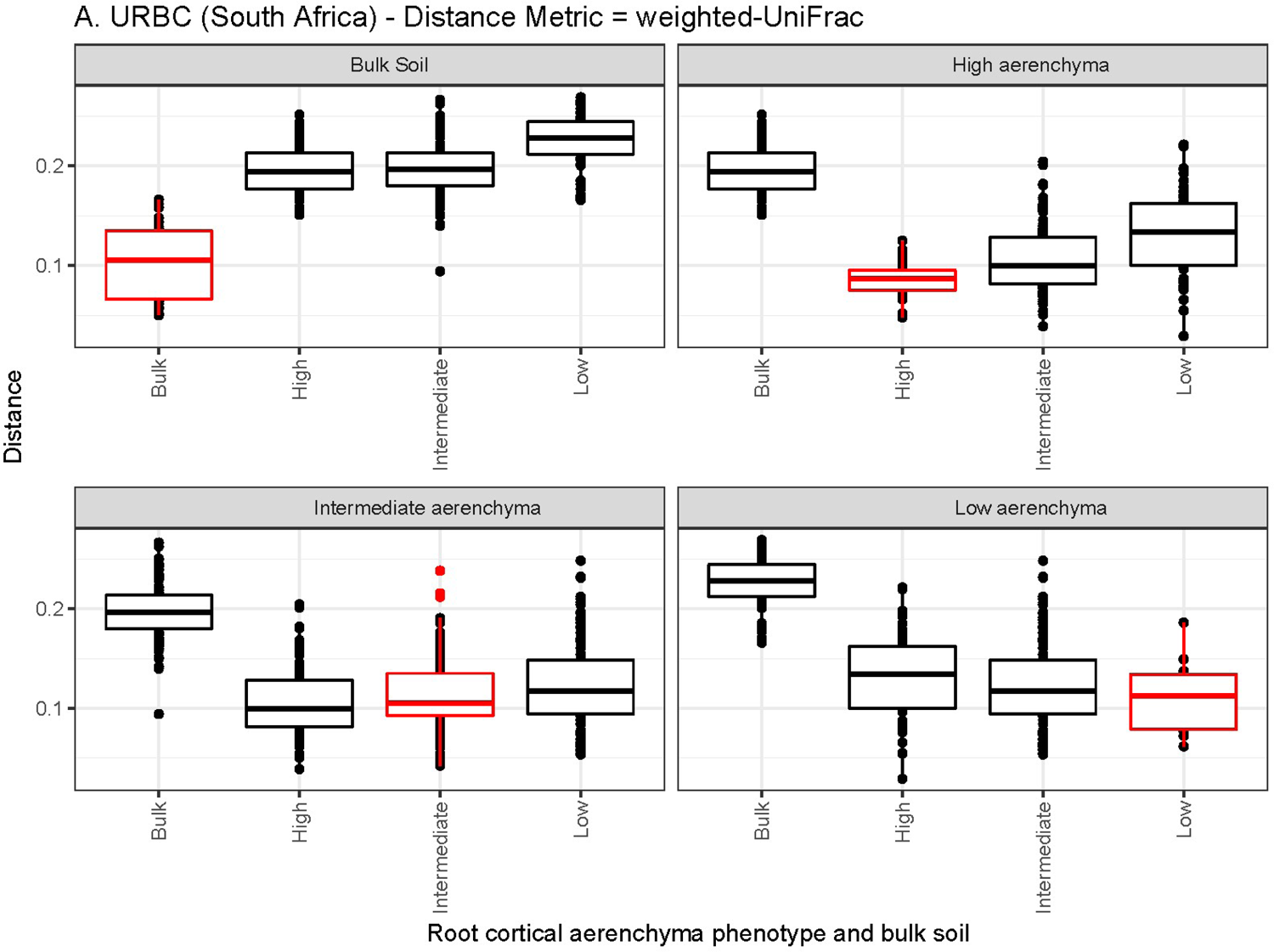
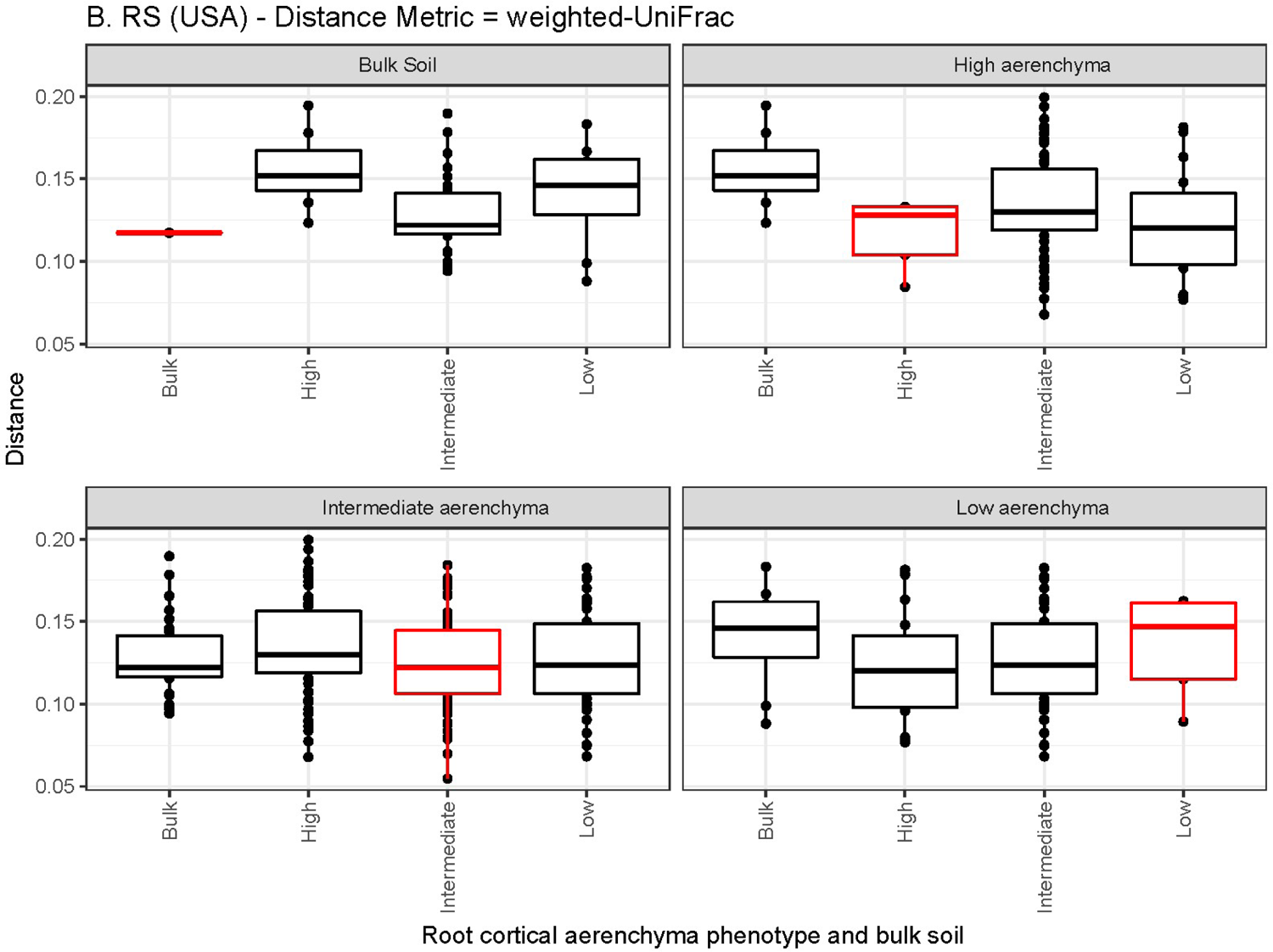
Boxplots comparing weighted-UniFrac distances between rhizosphere of plants of contrasting aerenchyma phenotype and bulk soil at URBC (A) and RS (B) on rarefied OTU counts. Horizontal box lines correspond to 25th, 50th, and 75th percentile; ranges are indicated by whiskers and points out of the boxes are outliers. Red-outlined boxplots correspond to self-comparison of the levels of phenotype or bulk soil.

**Supplementary Fig. 4.**
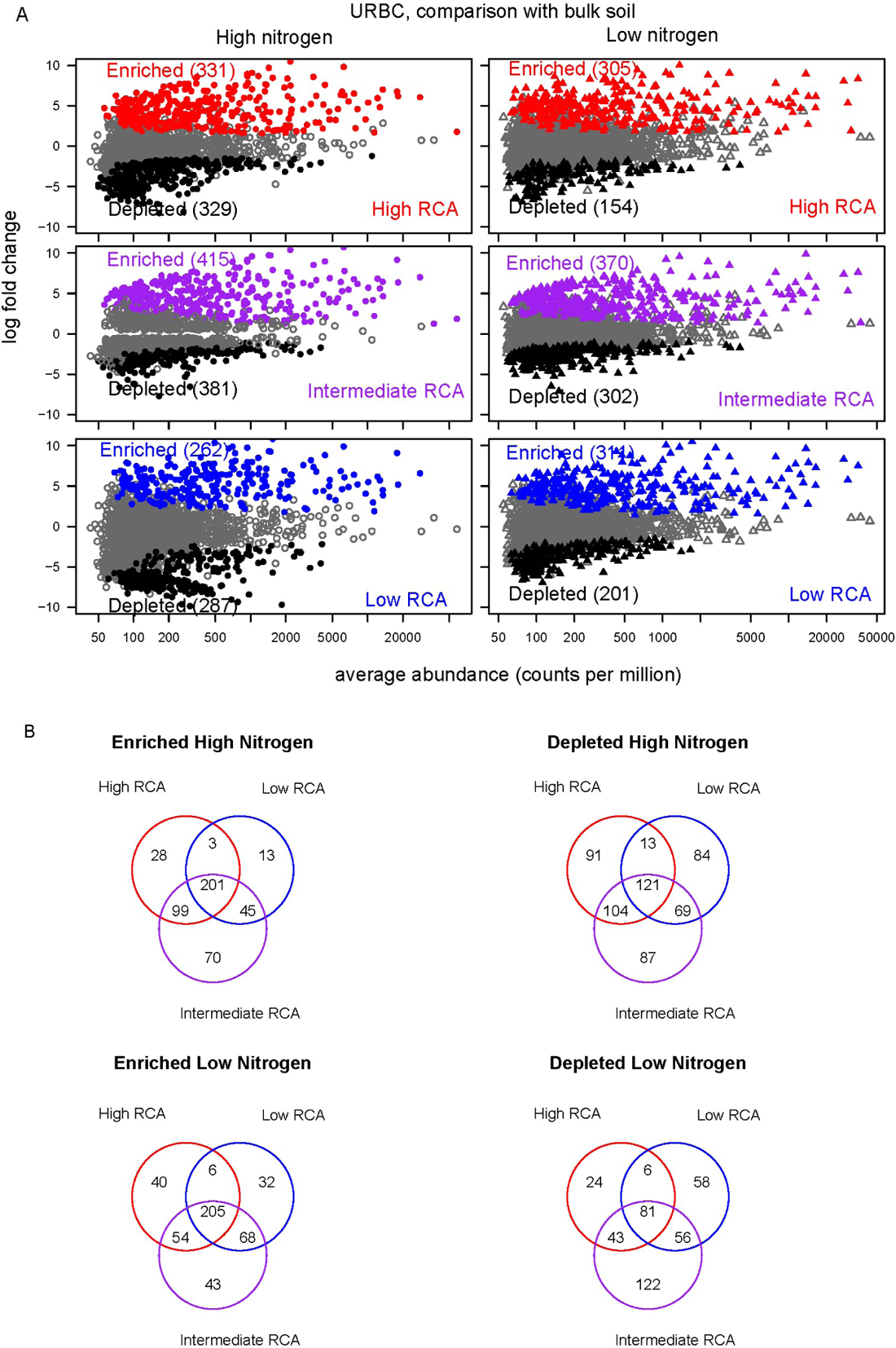
Abundance log change (y axis) of all the OTUs when rhizosphere (of each RCA rank) and bulk soil were compared at URBC (A). Colored points indicate differentially enriched (red, purple or blue) and depleted (black) OTUs according to a likelihood ratio test with *P*<0.01, and grey points were non-differentially abundant between the respective rhizosphere and bulk soil samples. Number of OTUs significantly enriched or decreased at each condition are in parenthesis. Number of the differentially enriched and depleted OTUs between each phenotype and bulk soil under the respective nitrogen level (B).

**Supplementary Fig. 5.**
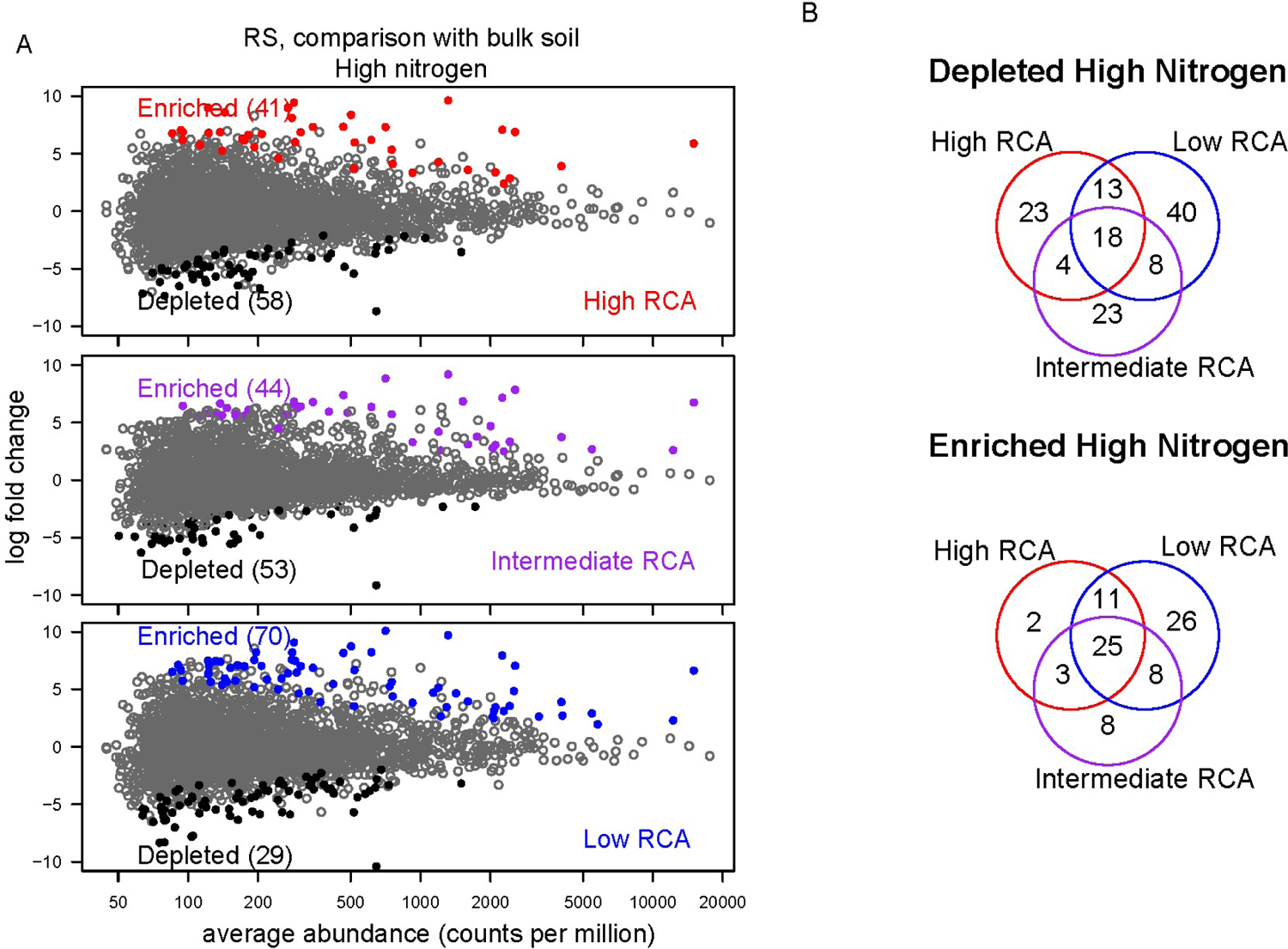
Abundance log change (y axis) of all the OTUs when rhizosphere (of each RCA rank) and bulk soil were compared at RS (A). Colored points indicate differentially enriched (red, purple or blue) and depleted (black) OTUs according to a likelihood ratio test with p<0.01, and grey points were non-differentially abundant between the respective rhizosphere and bulk soil samples. Number of OTUs significantly enriched or decreased at each condition are in parenthesis. Number of the differentially enriched and depleted OTUs between each phenotype and bulk soil under the respective nitrogen level (B).

**Supplementary Fig. 6.**
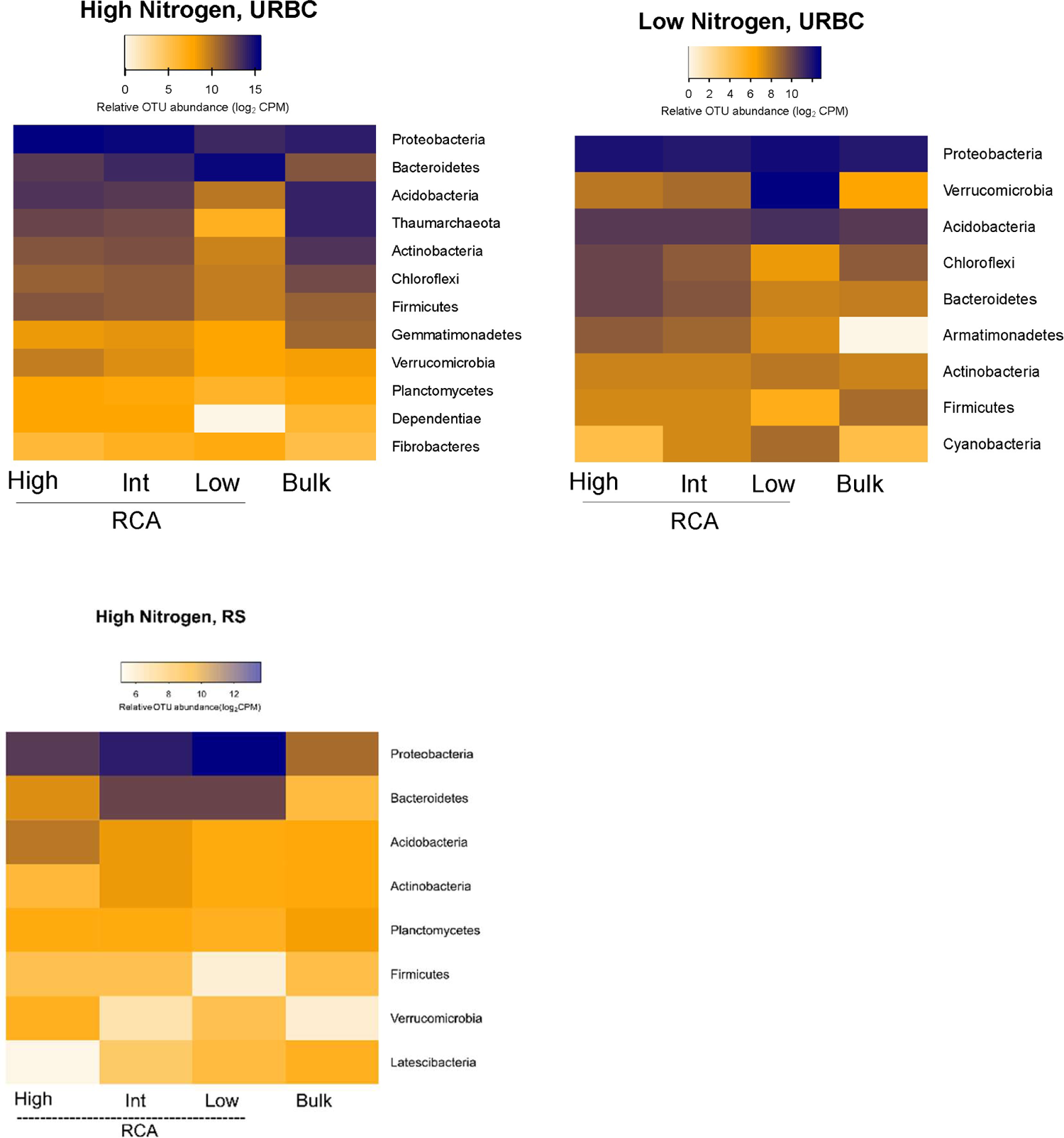
Mean relative abundances (counts per million, CPM; log2 scale) of RCA-sensitive OTUs (found as described in Fig. 6), summarized at phylum level under high and low nitrogen at URBC and under high nitrogen at RS and in comparison with the abundance values of bulk soil.

**Supplementary Fig. 7.**
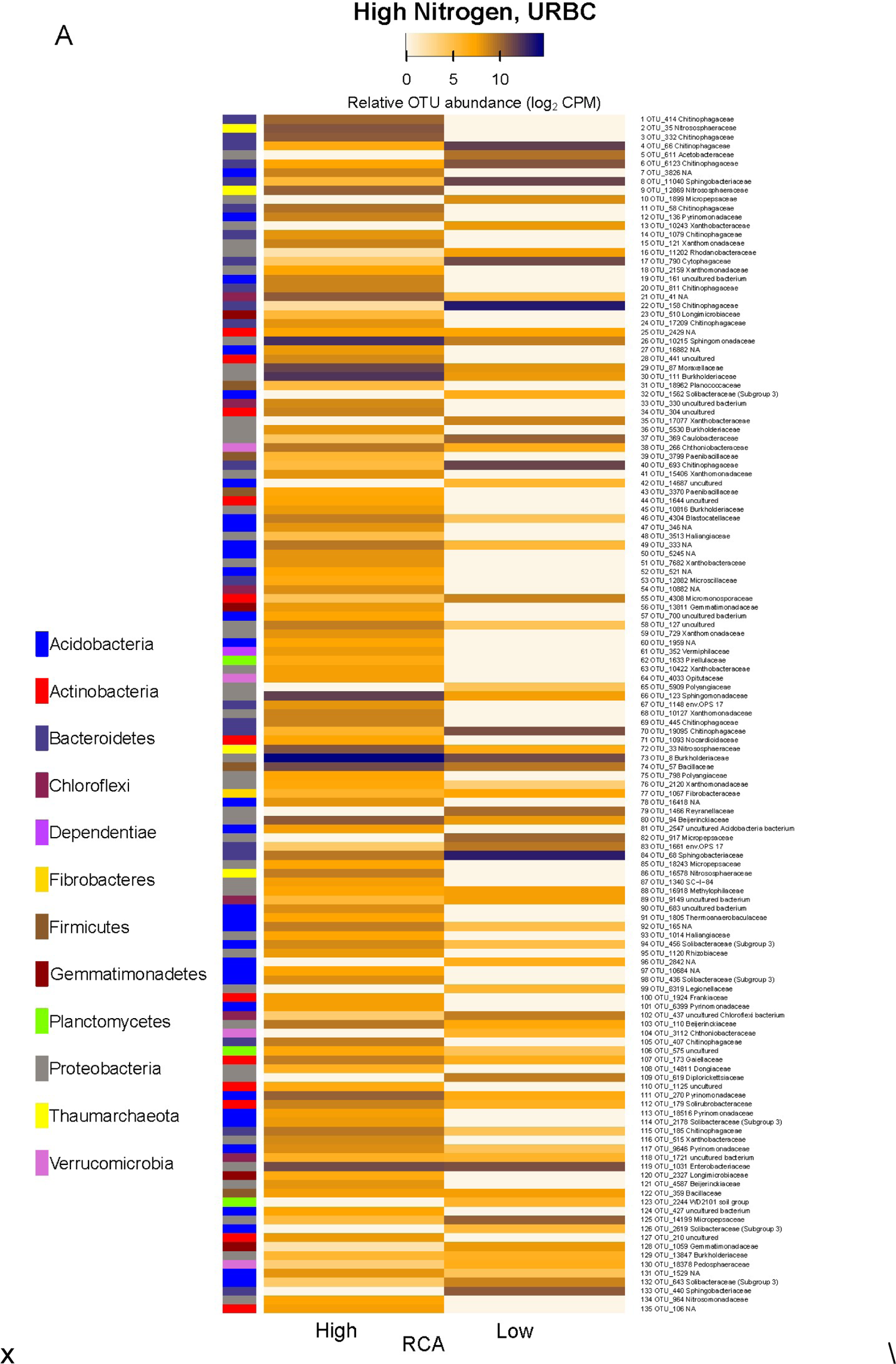
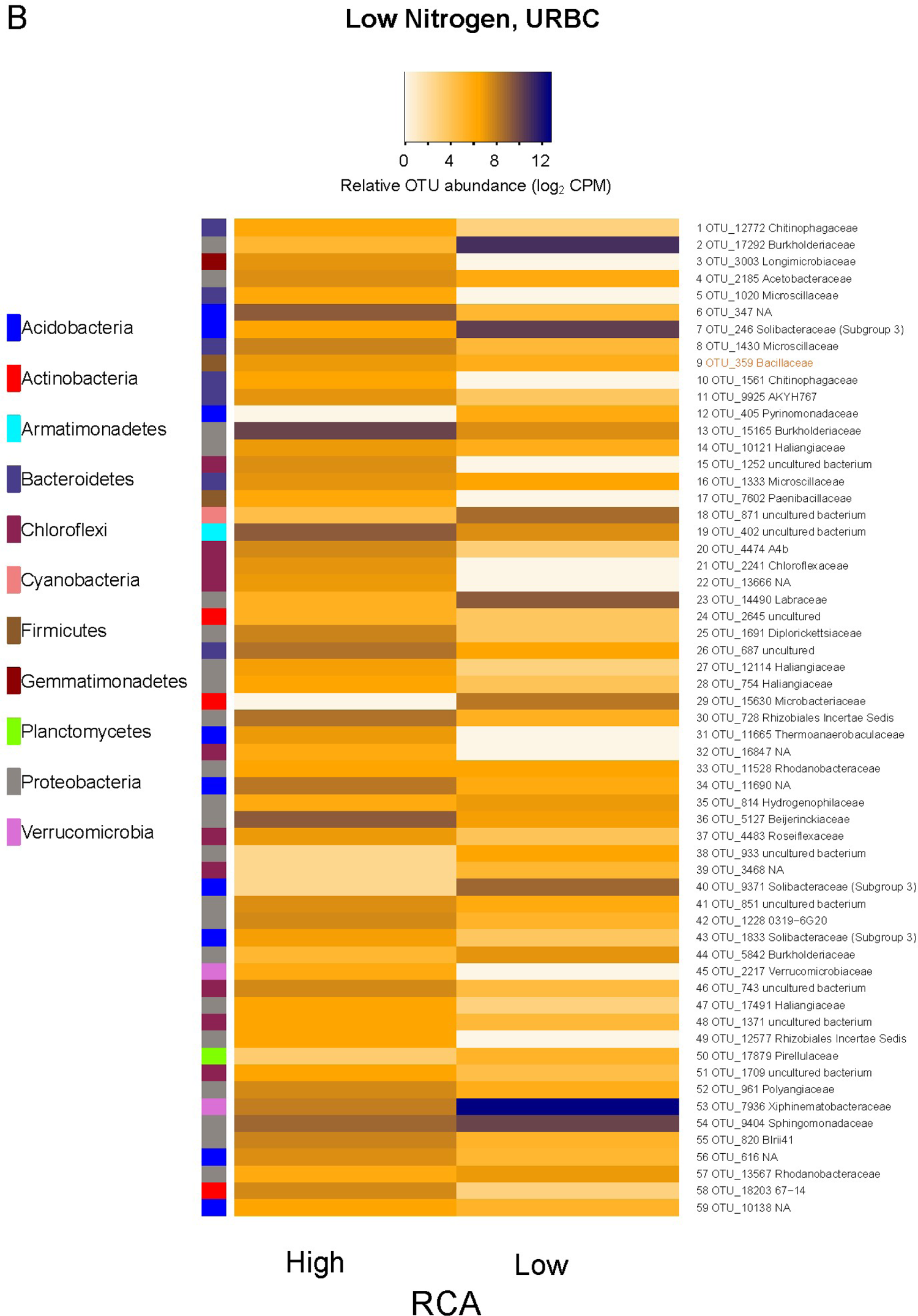
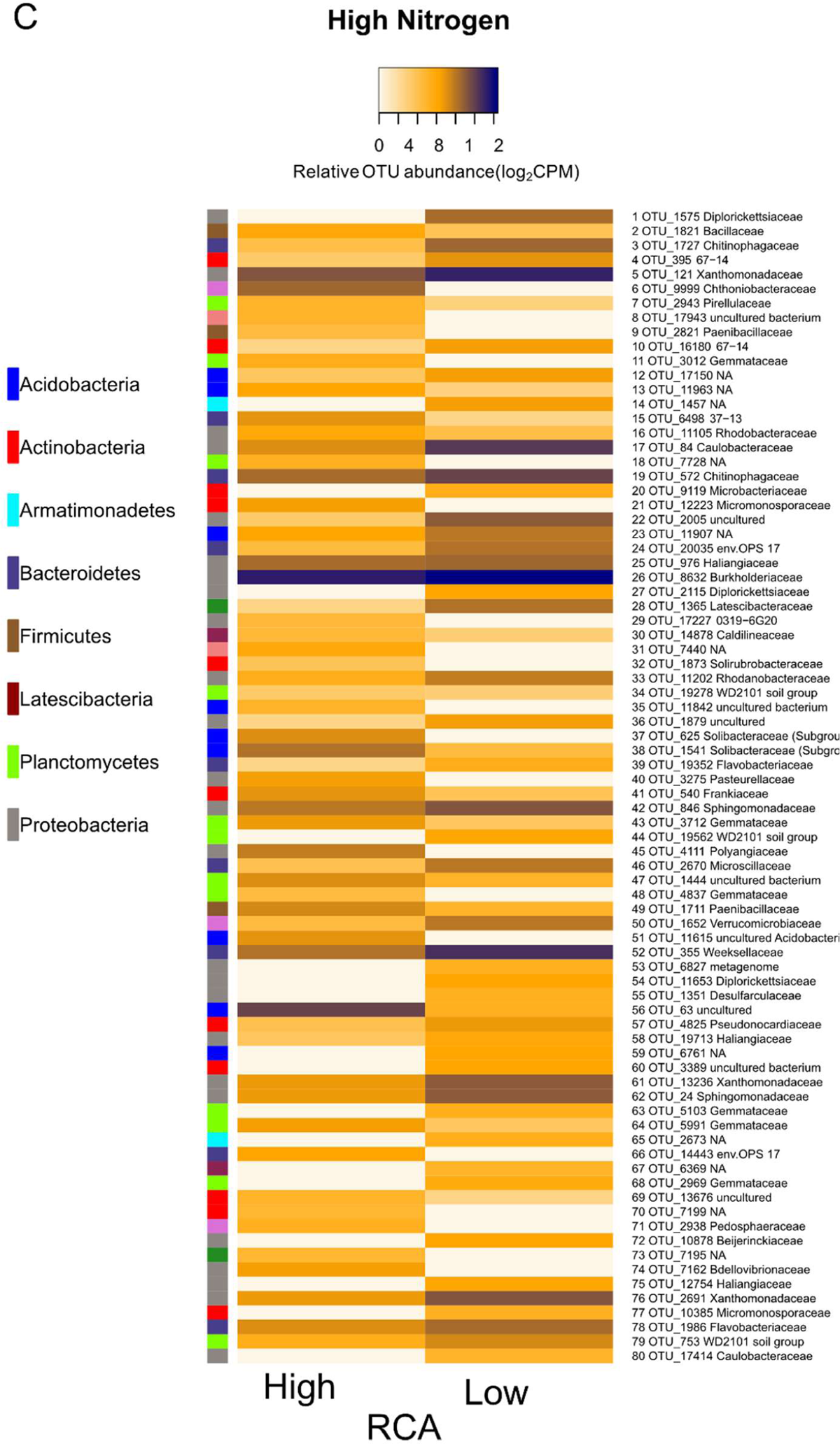
Mean relative abundances (counts per million, CPM; log2 scale) of RCA-sensitive OTUs (found as described in Fig. 6) at URBC (A,B) and at RS (C), summarized at family level (listed on right). Lists of the families and genera are also provided in File S1. The phyla are given in the colored bar of each graph (on left).

## Supplementary Results

### Taxonomy and putative functions of enriched OTUs at contrasting RCA phenotypes

Among the most abundant OTUs of high-RCA in high nitrogen conditions at URBC (South Africa), the families Comamonadaceae and Xanthomonadaceae and the genus *Sphingomonadaceae* (formerly classified as *Kaistobacter*, family Sphingomonadaceae) have been associated with disease-suppressive soils (Li et al., 2015; Liu et al., 2016). Bacteria of the nitrogen fixing families Beijerinckiaceae, Frankiaceae, and Rhizobiaceae where significantly enriched in high-RCA rhizospheres under high nitrogen, as well as ammonia oxidizing archaeans of the family Nitrososphaeraceae, and nitrate-reducers Gaiellaceae, Solirubrobacteraceae in addition to members of the phylum Acidobacteria and Actinobacteria that have been associated with the reduction of nitrate to nitrite by the assimilatory pathway (Albuquerque and da Costa, 2014a; Albuquerque and da Costa, 2014b; Campbell, 2014). Also, the family Chitinophagaceae which can degrade different organic macromolecues such as chitin and cellulose (McBride et al., 2014; Rosenberg, 2014), were found abundant in high-RCA plants under high nitrogen. At URBC, Low-RCA plants growing under high nitrogen conditions had three times less enriched OTUs compared to high-RCA. The two most abundant families among the enriched OTUs in low-RCA plants and high nitrogen were Sphingobacteriaceae and Chitinophagaceae (both from the phylum Bacteroidetes). Sphingobacteriaceae are aerobes chemoorganotrophs (Lambiase, 2014), and Chitinophagaceae are aerobes or facultative anaerobes with the potential to degrade macromolecules such as proteins, lipids, starch, pectin, chitin, carboxymethylcellulose or cellulose (McBride et al., 2014; Rosenberg, 2014).

Rhizospheres from low nitrogen plots at URBC had overall lower OTU abundances and a similar enriched-OTU distribution among the most abundant phyla compared to high nitrogen, with the difference that Armatimonadetes, a family generally considered aerobic of oligrotrophic metabolism (Lee et al., 2014) was uniquely enriched under low nitrogen in high and intermediate RCA plants, and Cyanobacteria was enriched in low-RCA plants (Supplementary Fig. 7). Another difference between the high and low nitrogen treatments at URBC was the lack of enrichment of the family Nitrososphaeraceae of high-RCA plants under low nitrogen. Similarly to the high nitrogen treatment, bacteria of the familiy Burkholderiaceae were enriched in high-RCA rhizospheres at low nitrogen, as well as nitrogen-fixing symbionts such as Rhizobiales and Beijerinckiaceae (Genus *Microvirga*, File S1), and the family Sphingomonadaceae from the phylum Proteobacteria that have been reported in soil and rhizosphere microbial surveys (Castillo et al., 2017; Lebeis et al., 2015; Schmid et al., 2017; Vik et al., 2013).

Among the OTUs with the highest abundance at high-RCA in high nitrogen at RS (USA), we found the genus *Oceanobacillus* (Phylum Firmicutes), previously reported in rhizosphere of halophyte plant species as betaine and proline producer and phosphate solubilizer (Mukhtar et al. 2018; El-Tarabily and Youssef, 2010). The most abundant OTUs enriched at low-RCA belonged to the familie Diplorickettsiaceae of the phyllym Proteobacteria (Genus *Aquicella*), a pathogen to protozoans (Albuquerque et al., 2018), and Chitinophagaceae of the phylum Bacteroidetes (genus *Chitinophaga*) common inhabitant of maize rhizospheres (Walters et al., 2018).

